# Vaccine waning and mumps re-emergence in the United States

**DOI:** 10.1101/185454

**Authors:** Joseph A. Lewnard, Yonatan H. Grad

## Abstract

Following decades of declining mumps incidence amid widespread vaccination, the United States and other high-income countries have experienced a resurgence in mumps cases over the last decade. Outbreaks affecting vaccinated individuals—and communities with high vaccine coverage—have prompted concerns about the effectiveness of the live attenuated vaccine currently in use: it is unclear if immune protection wanes, or if the vaccine protects inadequately against mumps virus lineages currently circulating. Synthesizing data from epidemiological studies, we estimate that vaccine-derived protection wanes at a timescale of 27 (95%CI: 16 to 51) years. After accounting for this waning, we identify no evidence of changes in vaccine effectiveness over time associated with the emergence of heterologous virus genotypes. Moreover, a mathematical model of mumps transmission validates our findings about the central role of vaccine waning in the re-emergence of cases: outbreaks from 2006 to the present among young adults, and outbreaks occurring in the late 1980s and early 1990s among adolescents, align with peaks in the susceptibility of these age groups attributable to loss of vaccine-derived protection. In contrast, evolution of mumps virus strains escaping pressure would be expected to cause a higher proportion of cases among children. Routine use of a third dose at age 18y, or booster dosing throughout adulthood, may enable mumps elimination and should be assessed in clinical trials.

**One Sentence Summary:** The estimated waning rate of vaccine-conferred immunity against mumps predicts observed changes in the age distribution of mumps cases in the United States since 1967.

## Introduction

Over the last decade, mumps outbreaks have thwarted the goal of eliminating indigenous mumps virus transmission in the United States by the year 2010 (*1*, *2*). Whereas over 90% of US-born children experienced mumps infections by age 20 in the pre-vaccine era (*3*), incidence declined substantially after licensure of a live attenuated vaccine (Jeryl Lynn strain) in 1967, in particular after the recommendation for its routine use among infants in 1977 as part of the measles-mumps-rubella (MMR) vaccine (*2*). Outbreaks among vaccinated middle school-and high school-aged children arose in the late 1980s, followed by sustained reductions in incidence after children were recommended to receive a second MMR dose at ages 4-6y (*4*). However, an ongoing resurgence in mumps cases began with a series of outbreaks on university campuses in 2006 (*2*). An older age of infection (ages 18-29y, compared to the pre-vaccine average of 5-9y) has been a defining feature these outbreaks (*5*), similar to recent experience in Canada, western Europe, and high-income Asian countries with routine MMR vaccination (*6*–*9*).

These circumstances are troubling on two fronts. First, as many as 10% of mumps infections acquired after puberty may cause severe complications including orchitis, meningitis, and deafness, in contrast to a milder clinical course in children that typically involves fever and parotid gland swelling (*10*). Second, a majority of mumps cases in recent outbreaks have been reported among young adults who received two vaccine doses as recommended (*11*). This observation has prompted concerns about suboptimal performance of the Jeryl Lynn vaccine currently in use (*12*).

It is unclear whether recent breakthrough outbreaks in vaccinated communities are due to waning of vaccine-derived immunity or to the emergence of mumps virus strains escaping vaccine-driven immunological pressure. Distinguishing between these possibilities is critical to policymakers and members of the scientific and medical communities: at issue is whether mumps can be eliminated by modifying vaccine dosing schedules, or if a new vaccine must instead be developed (*12*). To this end, we sought to distinguish waning of vaccine-derived protection from long-term changes in vaccine effectiveness (VE) against circulating mumps strains using data from previous studies. We then measured the potential impact of waning on population immunity over the decades since vaccine licensure, and used mathematical models to assess whether recent mumps virus transmission dynamics are more consistent with hypotheses of waning immunity or vaccine escape. We used these findings to evaluate alternative vaccination policies aiming to enhance protection among adults.

## Results

### Evidence of waning immunity in studies of vaccine effectiveness

Uncertainty about the protective efficacy of the Jeryl Lynn mumps vaccine—ranging from 95% following a single dose in randomized controlled trials (*13*) to <50% 2-dose effectiveness during recent outbreaks (*14*)—has undermined efforts to gauge population immunity. This variation in estimates of effectiveness permitted us to evaluate several hypotheses about the reasons mumps cases have re-emerged among vaccinated persons (*2*, *12*). Fitting a meta-regression model to data from prospective and retrospective cohort studies, we identified that the time elapsed since receipt of an individual’s last vaccine dose accounted for 66.4% of unexplained variation in published vaccine effectiveness estimates (**Figure 1A-C**). Applying our estimate of the vaccine waning rate to a model of exponentially-distributed durations of protection, we estimated that immunity persists, on average, 27.4 (95% confidence interval: 16.7 to 51.1) years after receipt of any dose. Among the 96.4% (94.0 to 97.8%) of individuals expected to mount primary responses to mumps vaccination, we thus expected 25% may lose protection within 7.9y (4.7 to 14.7y), 50% within 19.0y (11.2 to 35.4y), and 75% within 38.0y (22.4 to 70.8y).

The gradual replacement of predominantly genotype A mumps virus in the pre-vaccine era by other genotypes after vaccine introduction has also been suspected to contribute to diminished protection. However, we did not find evidence of a decline in vaccine effectiveness over calendar years, in particular after controlling for the effect of vaccine waning (**Figure 1D**). We also did not identify a difference in the duration of protection after first and second doses (**Figure 1E**). Whereas second doses were originally recommended to bolster immunity in case of failed “take” of the first dose, our findings suggest the second dose also restores immunity to levels achieved prior to waning of the first dose, thus extending protection to older ages. Taken together, our findings supported the central role of waning immune protection as a driver of variation in estimated effectiveness of mumps vaccine.

**Fig. 1.**
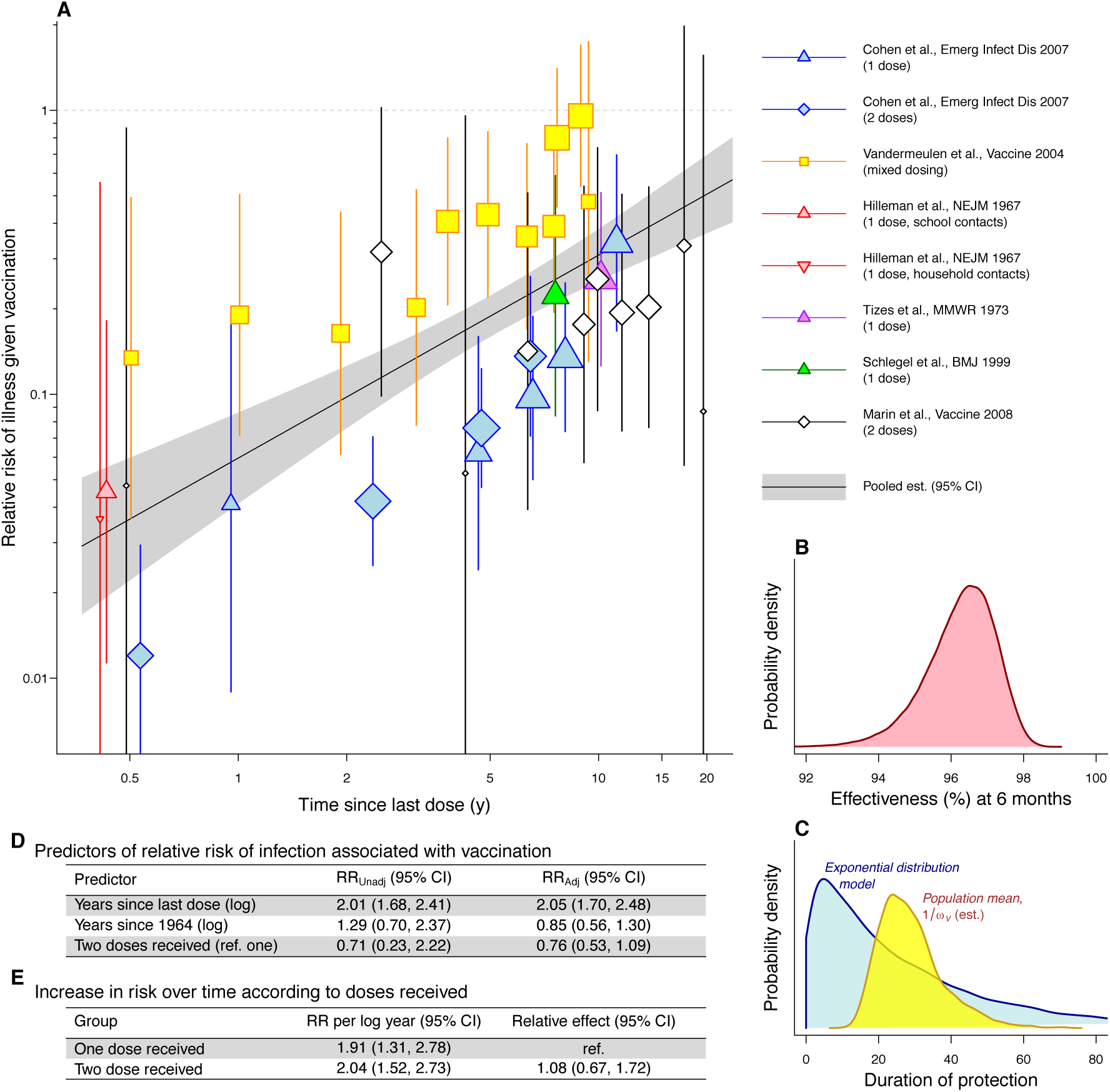
Synthesis of prospective and retrospective cohort studies estimating the relative risk of clinical mumps in vaccinated and unvaccinated individuals. (**A**) Estimates of VE (defined as one minus the relative risk of experiencing mumps given vaccination) have differed across studies; time since last dose accounts for 66.4% of residual variation in estimates after accounting for random sources of between-study heterogeneity. Points representing study-level estimates are scaled in size to reflect differences in sample size. (**B**) As of 6 months after receipt (the earliest time point assessed in primary studies), we estimated 96.4% (94.0 to 97.8%) of recipients are protected; we applied this as our estimate of the probability of vaccine “take”. (**C**) A parsimonious model of exponentially distributed durations of protection predicted loss of protection after, on average, 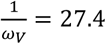 years (95%CI: 16.7-51.1), where *ω*_*V*_ is the waning rate, as indicated by the yellow plotted area. The blue plotted area illustrates the distribution of times to loss of protection for vaccinated individuals, generated by pooling exponential distributions parameterized using estimates of *ω*_*V*_. (**D**) Contrary to the hypothesis of reduced effectiveness against diverse mumps genotypes currently in circulation, we did not identify evidence of a decline in VE over time, whereas evidence of vaccine waning persisted in a model adjusting for calendar year. Unadjusted and adjusted estimates of the relative risk of clinical mumps given vaccination are calculated via meta-regression using data from the original studies. Adjusted relative risk estimates come from a model including all three covariates together. (**E**) Using this meta-regression framework, we identified no difference in waning rate after receipt of a first or second dose.

### Changes in population susceptibility to mumps virus after vaccine introduction

To understand the epidemiologic context of mumps resurgence in older age groups, we next sought to assess how waning vaccine-derived immunity and declining rates of natural transmission have impacted the susceptibility of the US population over the decades since vaccine introduction. We inferred the degree of immune protection as of 1967, when the vaccine was licensed, by fitting a mathematical model to reproduce epidemiological dynamics in the pre-vaccine era at steady state (*15*). We estimated that the basic reproductive number (*R*_0_) of mumps in the United States—the number of infections expected to result from an index case in a fully-susceptible population—was 4.79 prior to vaccine rollout, in agreement with previous estimates of 3–7 for high-income settings in the twentieth century (*16*, *17*). Allowing for loss of naturally acquired immunity did not improve model fit (*15*), consistent with longer-term persistence of high antibody titers after natural infection in comparison to vaccination in children (*18*).

The ongoing resurgence in mumps among young adults corresponds with cohort-specific changes in susceptibility resulting from vaccine waning and declining transmission over the decades since vaccine rollout (**Figure 2**). We estimated that 52.8% (41.6% to 63.1%) of adults ages 20-24y and 52.6% (42.4% to 61.3%) of adults ages 25-29y were susceptible to mumps virus infection in 2006 at the outset of the ongoing resurgence, in contrast to 33.8% (30.4% to 37.6%) and 25.2% (22.8% to 27.5%), respectively, as of 1990, and <10% in each age group before vaccine introduction. Susceptibility has also permeated older age groups amid the replacement of cohorts that experienced mumps as children. Whereas most individuals ages 65 and older had natural immunity as of 2016, we estimated that 29.2% (24.7% to 32.3%) of those ages 40-64y were at risk of infection. We expect these levels to continue increasing as transient vaccine-derived immunity supersedes previous infection as the main determinant of mumps susceptibility in the US population.

In a further validation of model predictions, the emergence and disappearance of mumps outbreaks among adolescents during the late 1980s and early 1990s corresponds to a transient increase in predicted susceptibility at ages 10-19y (**Figure 2**). We estimated that susceptibility at ages 10-14y peaked in 1991, when 45.8% (39.3% to 52.4%) of children in this age group were at risk for infection, as well as 43.0% (37.3% to 49.0%) of adolescents ages 15-19y. These levels reflect 2.85 (2.65 to 3.30)-fold and 3.96 (3.43 to 4.52)-fold increases in age-specific susceptibility, respectively, compared to the pre-vaccination era. Whereas breakthrough outbreaks beginning in the 1980s were hypothesized at that time to reflect inadequate responses of children to their first vaccine dose (*19*), our findings instead suggest that vaccine waning and declining natural exposure explain why adolescents were the population at highest risk for infection at that time.

We estimate that as of 2016, prevalence of susceptibility among children ages 10-14y declined to 34.8% (24.3 to 45.7%) due to the recommendation in 1989 for children to receive a second dose at ages 4-6y (*4*). Whereas most adolescents experiencing cases during the initial resurgence had received one dose of vaccine in keeping with the recommendations at that time (*20*, *21*), recent outbreaks have predominantly included individuals eligible to receive two doses (**see Supplementary Material**). Thus, the increasing age of infection in the US tracks with waning immunity after receipt of the second dose rather a continuation of cases within a single, under-immunized cohort.

**Fig. 2.**
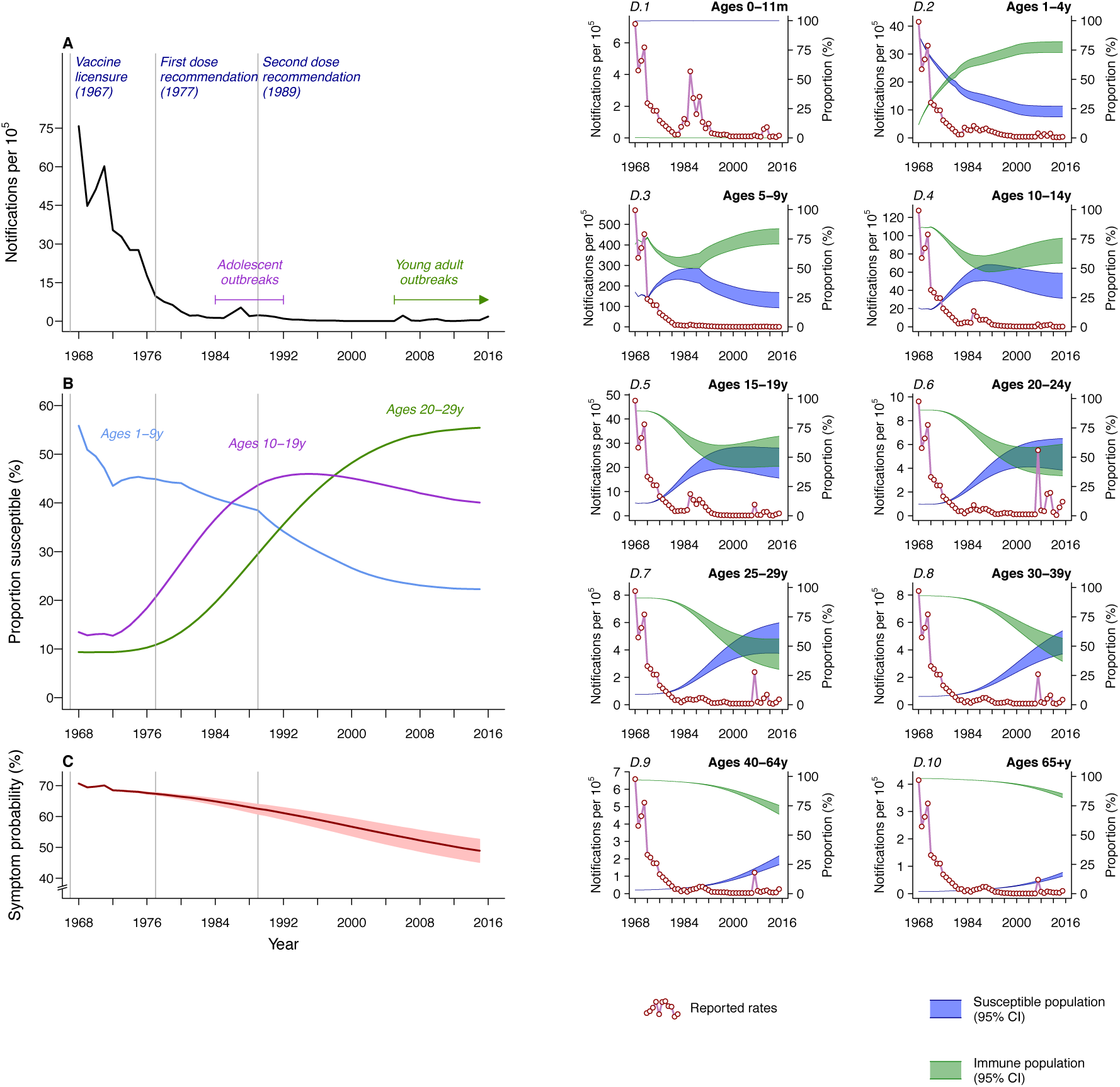
Mumps incidence and estimates of population susceptibility over time. (**A**) Overall notification rates declined following licensure in 1967, punctuated by outbreaks primarily among adolescents from 1984-1992 and recent outbreaks (2006 onward) centered among young adults. These outbreaks have corresponded with (**B**) peaks in the model-inferred proportion of individuals susceptible to mumps infection at ages 10-19y and 20-29y, respectively and (**C**) reductions in the proportion of infections that cause symptoms and are reported due to vaccine protection against symptoms. (**D**) Changes in the proportion of individuals susceptible to infection across different ages are plotted against case notification rates.

### Predicted transmission dynamics under vaccine waning and vaccine escape

Our analyses have suggested that reduced vaccine effectiveness relates primarily to waning protection rather than the emergence of mumps virus genotypes escaping vaccine-driven immunity. However, our ability to compare these hypotheses using data from previous studies is limited by a lack of data about genotype-specific protection. To better understand whether recent outbreaks are more consistent with vaccine waning or vaccine escape, we used a stochastic transmission model to compare expected epidemiologic dynamics under these scenarios in the year 2006, when the ongoing resurgence began. Using the approach taken above to update population immunity and transmission parameters in the absence of vaccine waning, we simulated the spread of mumps virus strains against which the vaccine provided partial protection in a population of 1 million. Strains capable of vaccine escape would be expected to cause higher-than-observed incidence among young children (**Figure 3A-E**): in contrast to a median age of 22y among cases reported in 2006, the predicted median age of cases approached 14.2y (8.3y to 21.7y) as strain-specific vaccine effectiveness declined to 0%. Although strains with lower ability to escape immune pressure may not concentrate to such an extent among children, our model predicted such strains would cause low incidence in a population unaffected by waning immunity (**Figure 3F**). Model-predicted overall rates of mumps incidence exceeded reported rates at lower degrees of cross-protection. Overall, model-predicted dynamics under vaccine waning provided a closer match to reported overall and age-specific incidence, with an expected median age of 22.3y (17.7y to 26.3y) among cases.

**Fig. 3.**
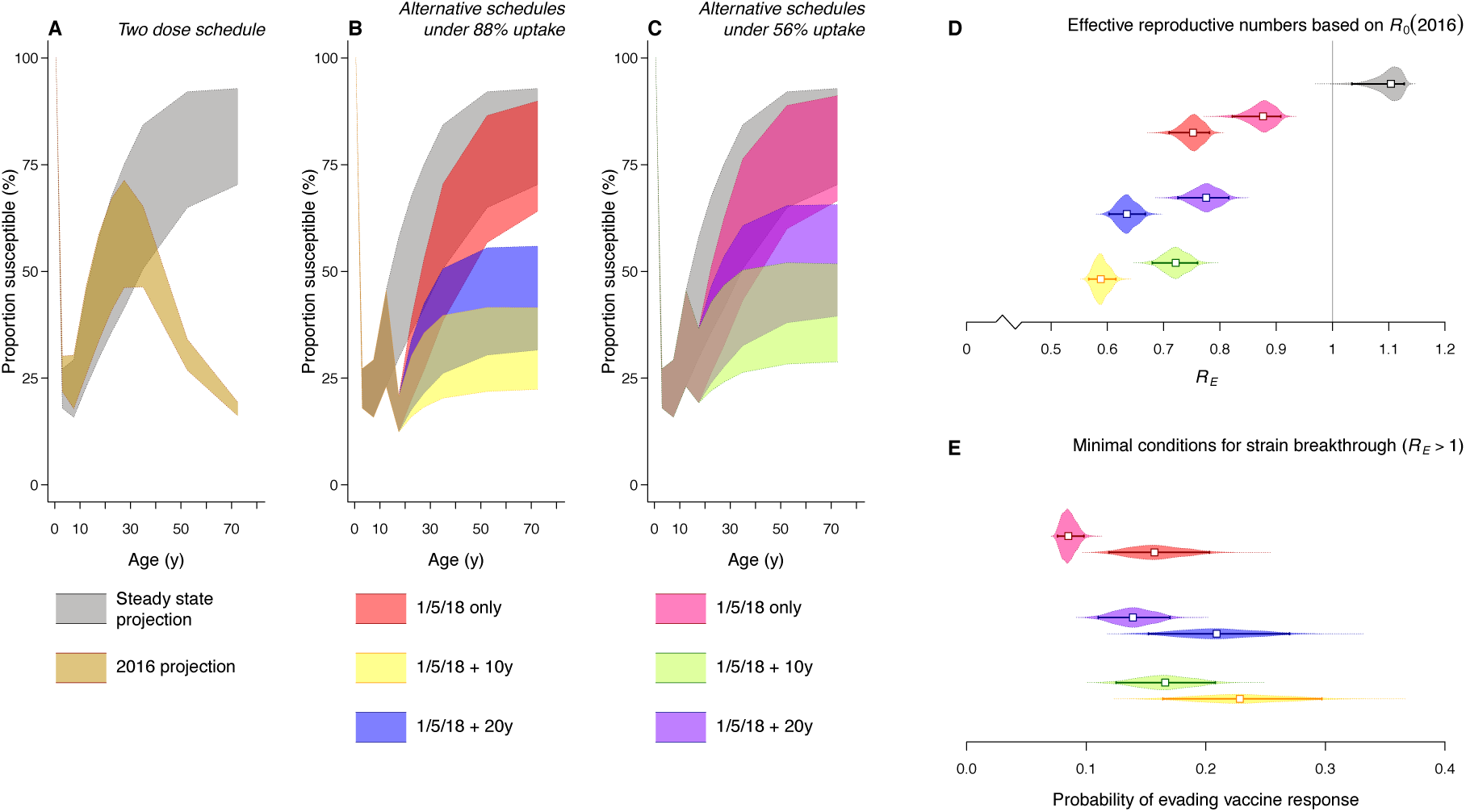
Anticipated transmission dynamics under scenarios of vaccine escape and vaccine waning. (**A**) A stochastic model of an emerging vaccine-escape strain of mumps virus in a vaccinated population predicts excess incidence in young age groups, in keeping with their higher burden of mumps historically. (**B)** In contrast, the fit of a model incorporating vaccine waning matches the observed age distribution. (**C, D**) Higher overall incidence rates and a younger age distribution of cases are predicted when immune responses to the vaccine offer minimal cross-protection against the circulating strain, as compared to the fit of the model with vaccine waning. (**E**) While the model with a vaccine-escape strain can reproduce the age distribution of cases at low degrees of immunological mismatch, (**F**) lower-than-reported incidence is expected under this scenario, again in contrast to the fit of a model with vaccine waning. Lines in (**E**) and (**F**) signify 95% confidence intervals.

### Potential impact of booster vaccination

If vaccine derived immunity wanes or confers shorter-lasting protection against genotypes currently in circulation as compared to those circulating in 1967, then administering additional vaccine doses may help control transmission by extending immune protection to older ages. Based on analyses of the effective reproductive number (*R*_*E*_), we found that immunity from two doses alone is unlikely to support elimination of endemic mumps virus transmission from the US in the long term. As birth cohorts exposed to high rates of transmission in the 20^th^ century are replaced by individuals whose protection comes only through vaccination, we expect *R*_*E*_ to approach 1.11 (1.04 to 1.13) (**Figure 4**). While administering a third dose by age 18y would not necessarily confer life-long protection based on our estimate of the time to loss of immunity, we nonetheless predict that this intervention could extend protection through young adulthood, thereby protecting age groups at risk in recent outbreaks. Low (56%) uptake of a third dose— matching adult compliance with recommended tetanus-diptheria toxoid booster doses—would be expected to sustain *R*_E_ around 0.88 (0.83 to 0.91), based on transmission dynamics in the US as of 2016. Under a more optimistic scenario of 88% third-dose coverage—where third-dose uptake equates second-dose uptake among already-immunized individuals—we expect *R*_*E*_ to approach 0.77 (0.72 to 0.79) as cohorts previously exposed to high transmission rates age out of the population.

Whereas we estimate most older adults are currently immune to mumps virus due to previous infection, our modeling suggests neither a two-dose nor three-dose vaccination program would be expected to protect over 50% of adults beyond the age of 40y in the long term. This concern may motivate the use of routine booster doses in adulthood (**Figure 4**). Based on our model, we expect that administering additional doses every 10y or 20y would lead to sustained protection in, at minimum, 68.0% (58.5% to 77.6%) and 55.2% (44.1% to 68.4%) of the population, respectively, under a scenario of 88% vaccine coverage; at the lower (56%) coverage level, we estimated protection among, at minimum, 59.0% (48.2% to 71.2%) and 45.5% (34.3% to 60.5%) of adults with dosing every 10y or 20y, respectively. Maintaining high levels of immunity in the population through repeated dosing may also help to contain emergence of novel mumps virus strains (**Figure 4E**). To sustain *R*_E_≥1 under three-dose schedules, we estimated that an emerging strain would require, at minimum, 8.5% (7.6% to 9.8%) to 15.7% (11.9% to 20.3%) probability of causing infection in exposed persons otherwise protected by vaccination. Adding 10y booster doses increase this threshold probability to between 16.6% (12.5% to 20.8%) and 22.9% (16.4% to 29.7%) at varying levels of vaccine coverage.

**Fig. 4.**
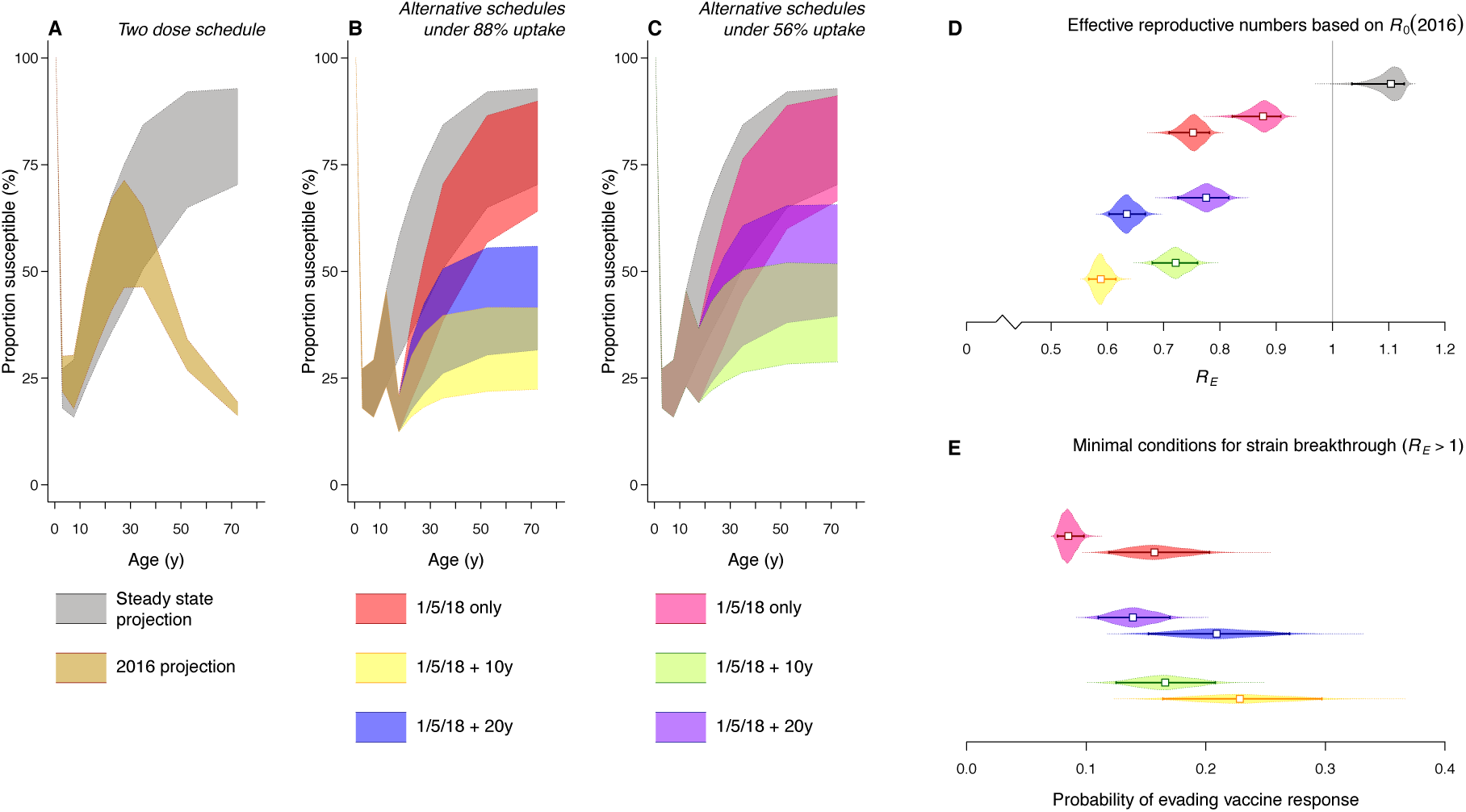
Age-specific immunity and transmission dynamics under two-and three-dose vaccine schedules. **(A)** Cohorts over 40 years of age as of 2016 were exposed to endemic transmission prior to and shortly after vaccine rollout and likely retain life-long protection. However, a population protected only by 2-dose vaccination would be expected to experience high levels of susceptibility over age 20y. (**B** and **C**) Our modeling suggests the duration of protection can be extended through young adulthood by adding a third dose around age 18, whereas routine booster doses every 10y or 20y would be expected to sustain longer-term protection. Lines and shaded areas delineate median estimates and 95% confidence intervals, respectively. (**D**) Under transmission dynamics estimated as of 2016, protection in young adult age groups achieved through the use of a third dose is expected to reduce the effective reproductive number (*R*_*E*_) below 1, whereas we predict *R*_*E*_ to approach 1.10 under the two-dose schedule as cohorts that experienced high rates of mumps infection age out of the population; larger reductions in *R*_*E*_ are sustained at higher levels of coverage and with more frequent dosing. (**E**) In turn, these levels of protection translate to a stronger barrier against emergence of vaccine-escape strains. Whereas a novel strain with 8.5% (7.6% to 9.8%) probability of evading vaccine-induced immunity and infecting a vaccine-protected individual would be expected to succeed under a three-dose schedule with low coverage, we estimate that a novel strain would require 22.9% (16.4% to 29.7%) probability of infecting such an individual to emerge in a population with 88% uptake of the third dose and 10y boosters. Lines denote 95% confidence intervals and shaded areas represent distributions around *R*_*E*_ estimates.

## Discussion

Resurgent outbreaks centered among young adults have brought renewed attention to mumps following decades of progress toward its elimination from the US (*2*, *12*). Understanding why cases have re-emerged is essential for determining how to contain the disease through vaccination. Our analyses show that vaccine derived immune protection wanes over time, provide estimates of the waning rate, and demonstrate that this waning immunity accounts for susceptibility in the age groups experiencing outbreaks over the decades since vaccine introduction in the US. In contrast, changes in the circulating genotypes of mumps virus over this same period have not been associated with reductions in vaccine effectiveness; moreover, our modeling suggests vaccine-escape mumps virus strains would be expected to cause disproportionate incidence among younger children. Guided by these findings, we identify that routine use of a third dose vaccine dose around age 18y, with or without regular dosing in adulthood, could help maintain immune protection in the population.

Distinguishing between the contributions of vaccine waning and the emergence of vaccine-escape virus strains to mumps resurgence helps to inform whether novel vaccines are needed to control transmission (*22*, *23*). Our findings that vaccine effectiveness has not declined amid the replacement of genotype A mumps viruses (from which the Jeryl Lynn vaccine strain was derived), and that the age distribution of recent cases is inconsistent with expectations under vaccine escape, are in agreement with several lines of evidence that mumps vaccination protects broadly against heterologous strains (*24*). Neutralizing antibody responses to the Jeryl Lynn strain are effective *in vitro* against wild-type mumps virus strains responsible for recent outbreaks among vaccinated individuals (*25*, *26*), and genetic distinctions have not been identified between strains isolated from vaccinated and unvaccinated mumps patients (*27*). Similar, high levels of efficacy and effectiveness have been estimated for the Jeryl Lynn and Urabe (genotype B-derived) vaccines, further substantiating the notion of cross-neutralizing or monotypic immune responses (*12*). Nonetheless, epidemiological studies of outbreaks caused by distinct virus lineages, and in populations exposed to different circulating mumps viruses, can better characterize genotype specificity in the strength or duration of vaccine protection.

Although the clinical efficacy of a third dose has not been assessed in a trial, several observations suggest effectiveness of extended vaccine schedules. First, whereas congregated US military populations resemble high-risk groups based on their age distribution and close-contact environments, no outbreaks have been reported in the military since a policy was adopted in 1991 of administering an MMR dose to incoming recruits, regardless of vaccination history (*28*). Second, in limited observational studies of vaccine campaigns undertaken in response to recent outbreaks, third-dose recipients have tended to experience lower incidence rates than non-recipients (*29*–*32*. However, trials demonstrating the clinical effectiveness of adult vaccine doses are needed to guide policy. Third-dose campaigns undertaken at the tail end of outbreaks have not been designed optimally to measure vaccine effectiveness (*29*, *31*), and conflicting evidence about immune responses to third doses in adulthood has proven difficult to interpret without known immunological correlates of protection (*33*–*35*. If a third dose at age 18y does not confer adequate protection, modifying MMR vaccines to improve the magnitude or duration of immune responses against mumps virus may present a viable alternative. Notably, the mumps component of the MMR vaccine induces lower-avidity antibody responses, and weaker specific memory B-cell proliferation, than the measles and rubella components (*36*, *37*).

Our findings support previous observations of waning vaccine-derived immunity against mumps virus (*38*) and are validated by consistency between the age distribution of reported cases and model-predicted susceptibility over time. Several limitations of the analysis should be considered. Our use of aggregated rather than individual-level data from vaccine effectiveness studies contributed to an imprecise estimate of the time to loss of immunity, in turn limiting the precision of our estimates of population susceptibility. Individual-level data from post-licensure vaccine studies could support better inferences about the magnitude and duration of vaccine protection, thus aiding policy decisions. Identifying immunological correlates of protection from such datasets would also aid evaluations of alternative vaccination schedules and measurements of population immunity. Last, our analysis addresses mumps epidemiology in the United States, where levels of immunity within particular birth cohorts may differ from levels in settings that introduced routine mumps vaccination later or use different vaccine schedules. The burden of cases and prevalence of immunity across ages or birth cohorts should be considered to guide vaccination policy within specific countries.

Analyzing nationally-aggregated incidence datasets also limited our ability to investigate how geographic or socioeconomic differences in vaccine uptake and contact rates contribute to the dynamics of focal outbreaks, as might occur in close-contact settings such as university dormitories (*39*). However, our inferences about vaccine waning and the changing age distribution of mumps cases offer insight into why mumps resurgence has been possible throughout geographically and socioeconomically distinct communities. In this regard, the widespread re-emergence of mumps in vaccine-compliant communities stands in stark contrast to the focal re-emergence of measles in communities with low vaccine coverage (*11*).

Changes in the epidemiology of mumps have implications for disease surveillance. Diminished clinical awareness of mumps, expectations that it appears in pediatric rather than adult populations, and protection against symptoms in vaccinated individuals (*40*) may limit routine detection of cases, and thus bias disease reporting. Indeed, serological surveys have provided evidence of higher-than-reported rates of mumps virus infection in the US prior to 2006 (*28*, *41*). The tendency to identify outbreak-associated cases through contact tracing may also favor detection of cases in university campuses and other closely-connected populations, underscoring the importance of serosurveys to assess the extent of transmission in the community. Serological datasets can also help establish whether lower rates of immunological “boosting” through natural exposure, which our analysis does not address, have contributed to population susceptibility.

The ongoing resurgence in mumps among young adults has undermined previous enthusiasm about near-term elimination of this disease from the US (*1*). Our analysis suggests that vaccinated individuals lose protection against infection on average 27 years after receipt of their last dose, and that this rate of vaccine waning explains susceptibility in adolescent and young-adult cohorts at the time of post-licensure outbreaks in these age groups. Re-emergence of mumps among older, previously-vaccinated individuals whose immunity has waned parallels recent experience with varicella outbreaks affecting immunized communities as a result of waning vaccine-derived protection (*42*). As demonstrated in mumps epidemiology, immunity in previously infected cohorts may buffer transmission and delay breakthrough epidemics from occurring until decades after vaccine introduction. These observations indicate the need for either innovative trial designs to measure the benefit of extending vaccine dosing schedules or novel vaccines to address the problem of waning vaccine-induced protection (*43*).

## Materials and Methods

Full technical details pertaining to our analyses are presented in the Supplementary Materials; a brief summary is provided here.

### Meta-analysis of vaccine effectiveness studies

We performed a systematic review of prospective and retrospective cohort studies calculating effectiveness of the Jeryl Lynn-strain mumps vaccine via a PubMed search and citation tracking. We used an inverse variance-weighted meta-regression model, accounting for study-level heterogeneity, to measure unadjusted and multivariate-adjusted associations of the following variables with study-level estimates of the relative risk of infection associated with vaccination:

1. Time since receipt of the last vaccine dose, indicating vaccine waning;
2. Time from 1964 (when the Jeryl Lynn vaccine was developed) to the year of mumps exposure, indicating long-term changes in vaccine effectiveness associated with changes in circulating genotypes; and
3. Vaccine doses received, interacted with time since last dose to test for differential waning of first and second doses.

Regression model summary statistics indicated the proportion of variance explained by these covariates. We used our estimate of the association between instantaneous risk of infection and time since vaccination to fit an exponential distribution to the duration of vaccine derived immune protection, and used this fitted distribution as the basis for further modeling.

### Modeling population immunity and mumps virus transmission

We used a system of ordinary differential equations to describe changes in the population of susceptible and immune persons, partitioned into those who had and had not received mumps vaccine doses based on reported vaccine coverage. We back-calculated changes in natural immunity in the population based on the relation

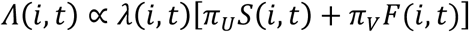

between reported incidence rates (A) in age group *i* and year *t* and the force of infection to which individuals were exposed (λ), the populations of susceptible unvaccinated *(S)* and vaccinated *(F)* persons, and the probabilities (*π_U_* and *π_V_*, respectively) for these individuals to experience symptoms if infected. Age-specific incidence reports were collated from nationwide surveillance (*44*). We inferred starting levels of immunity in the population (as of 1967) by fitting a mathematical model of mumps transmission to recapitulate age-specific incidence in the pre-vaccine era at steady state. We implemented stochastic realizations of an extended version of the model to compare predicted incidence during an introduced outbreak against observations from the year 2006, under scenarios where we assumed waning of immunity or circulation of strains with differential risk of infecting vaccinated persons.

### Assessing extended-dose strategies

We compared the long-term performance of different vaccination schedules including the addition of a third dose by age 18y and the use of routine boosters at 10y or 20y intervals through adulthood. We calculated the prevalence of age-specific immune protection achieved under these strategies and resulting values of the effective reproductive number (*R_E_*), describing the number of cases an infectious individual would be expected to cause under prevailing conditions. We also calculated the minimum probability of immune escape a novel mumps virus strain would need to invade a population protected under these different strategies, defined as the minimum probability of infecting a vaccinated, protected individual upon exposure such that *R*_*E*_≥1.

## Acknowledgments

The authors thank Lucy Li, Sarah Cobey, Pardis Sabeti, Shirlee Wohl, Nathan Yozwiak, Greg Armstrong, and Marc Lipsitch for input.

## Author contributions

JAL and YHG conceived of the study, JAL performed the analyses with input and discussion from YHG, and JAL drafted and YHG edited the manuscript.

## Data and materials availability

Code for replicating analyses and figures is available at github.com/joelewnard/mumps.

## Supplementary Materials

### SM1. Meta-analytic estimation of vaccine waning rates

#### SM1.1. Literature search

We performed a systematic literature search to identify studies from which we could infer vaccine waning. We searched the Boolean phrase mumps AND vaccin* AND ((second* AND fail*) OR wan*) in PubMed (obtaining 77 results) and Google Scholar (considering the first 300 results). We also aggregated all references listed in three recent articles (*9*, *12*, *14*) reviewing studies of mumps vaccine performance (totaling 234 results), and all citations of these three articles in Google Scholar (totaling 370 results as of 6 June 2017). We similarly performed forward-and back citation tracking of included studies. We included articles:

1. written in English;
2. presenting results of prospective or retrospective cohort studies of mumps vaccination and disease incidence, and thereby enabling calculation of the relative risk of laboratory-confirmed clinical mumps according to vaccination status;
3. clearly indicating the time elapsed between receipt of the last vaccine dose and onset of exposure to mumps transmission; and
4. (if individuals in the study were born within five years after the introduction of routine mumps vaccination): excluding individuals with serologic or clinical history of mumps infection, thus preventing misclassification bias that could otherwise be confused for evidence of vaccine waning (*45*). We assumed low risk of previous mumps infection for individuals born more than five years after implementation of routine vaccination.

Two studies (*46*, *47*) which met the conditions listed above were excluded from the analysis because they lacked an unvaccinated reference population, preventing the estimation of effect sizes. The included studies are listed in Table S1.

**Table S1:**
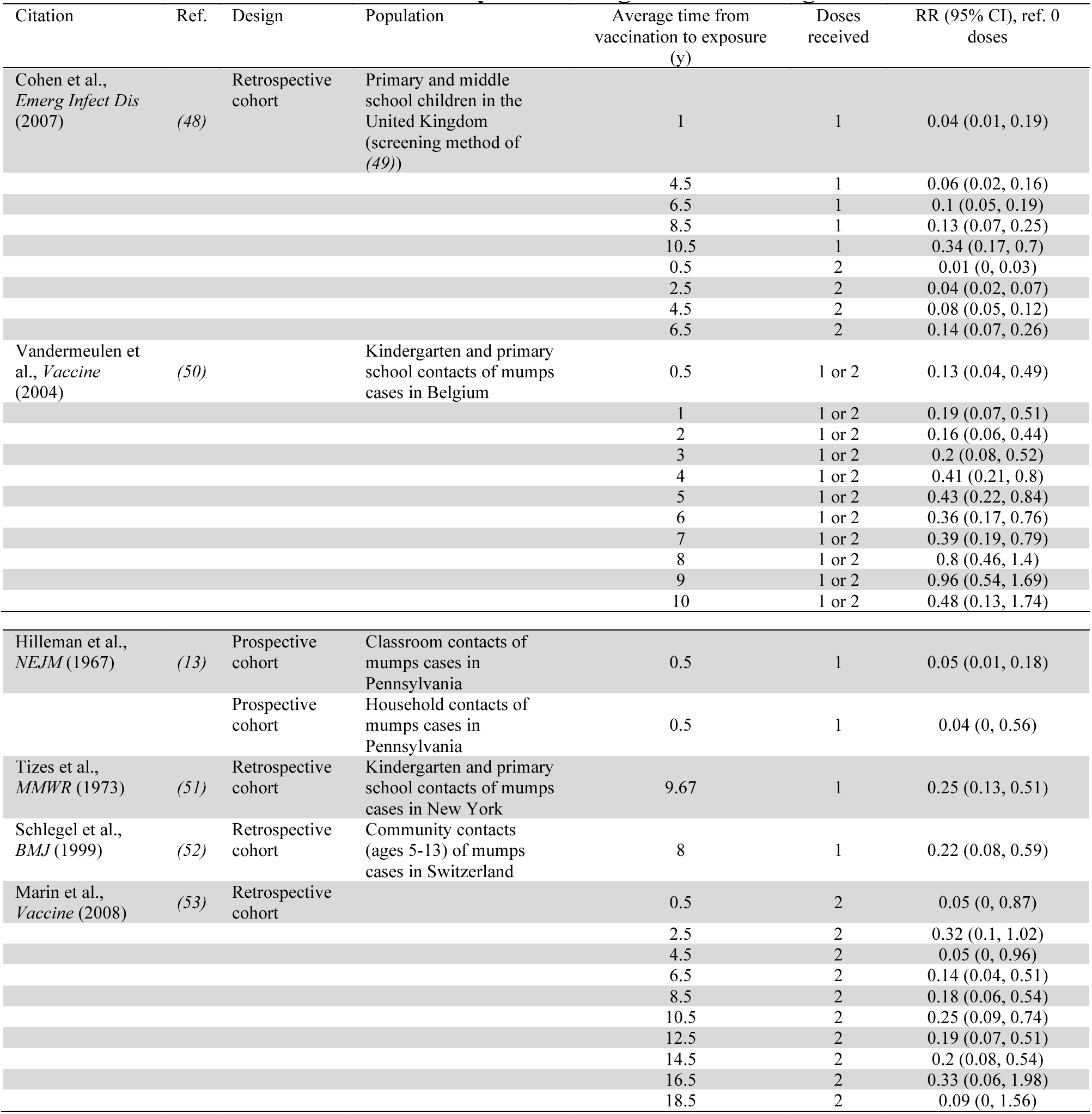
Studies included in meta-analysis assessing vaccine waning.

We obtained the number of mumps cases reported among total vaccinated and unvaccinated study populations together with vaccination status and time from vaccination to exposure. When possible, we extracted subsamples from individual studies wherein subjects differed in time since vaccination.

#### SM1.2. Testing for differences in protection with time since vaccination

We fitted an inverse variance-weighted model of log-transformed relative risk estimates against time since vaccination, allowing for fixed study-level effects based on a comparison of the Bayesian Information Criterion from fixed-and random-effects models (BIC_FE_=69.6, BIC_RE_=73.6). By this same measure, the model including time since receipt of the last dose provided a better fit to data than a null model with study-level intercepts alone (BIC_null_=103.1). We measured the proportion of residual variance from the null model (*M*^0^) explained by the model including time since last dose (*M*^*^) as the reduction in weighted squared errors:

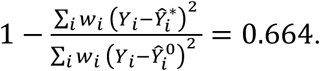

Weights *w*_*i*_ were proportional to the inverse of the variance of estimates *Y*_*i*_ from original studies. Estimated from the pseudo-*R*^2^ value, the model including time since last dose and study-level intercepts explained 88.9% of variation in previous estimates of effectiveness. We used the same approach to test for a significant association between the year of exposure among cases included in the study and time since 1964, which we expected to indicate any long-term change in vaccine effectiveness associated with changes in circulating genotypes of mumps virus; however, we identified no significant effect.

We next tested whether waning rates differed for children who had received one or two doses of vaccine. Excluding one study which presented aggregated data from 1-dose and 2-dose recipients (*50*), we identified no improvement in model fit when allowing for differential rates of loss of protection among 1-dose and 2-dose recipients (BIC_Any-dose_=56.1, BIC_Dose-specific_=58.3). Moreover, the 95% confidence interval around the estimated fold change in loss rates after a second dose (ref a single dose) spanned 1, indicating no significant difference in waning rate.

#### SM1.3. Estimating rates of waning of vaccine-derived protection

Our analysis of aggregated data from studies did not allow us to infer the distribution of waning rates across individuals. As a result, assuming exponentially-distributed time to loss of protection provided the most parsimonious basis for predicting for the proportion of vaccinated individuals who, conditioned upon initial vaccine “take”, would retain protection at a given time. We further assumed our estimate of vaccine effectiveness at 6 months from the above regression model approximated the proportion of doses initially conferring protection (v), as 6 months was the shortest average duration of follow-up from time of vaccination among included studies (**Table S1**). Our estimate of vaccine effectiveness at this time point was 96.4% (94.0-97.8%).

We fitted an exponential waning rate *ω*_*V*_ by minimizing the sum of squared errors between:

1. the proportion of individuals initially protected by the vaccine who retain protection under exponentially-distributed durations of protection, exp(-*ω*_*V*_*t*)/*v*, where *t* indicates years since receipt of the last dose (*t*>0.5), and
2. estimates of the ratio of vaccine effectiveness at time *t* to time *t*=0.5 under the meta-regression model:

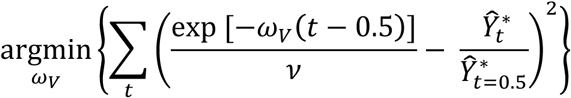

for *t*={0.5, 0.51, 0.52, …, 18.5}, corresponding to the range of observations included in the original estimates (**Table S1**). To propagate uncertainty from the initial regression estimates, we constructed the distribution of *ω*_*V*_ by fitting against independent samples of {*v*, **Ŷ**^*^} obtained from the multivariate-normal distribution of regression parameters.

### SM2. Inferring transmission dynamics in the pre-vaccination era

#### SM2.1 Model of mumps transmission

We modeled steady-state transmission dynamics of mumps in the US prior to vaccine introduction (1967) to estimate starting-time, age-specific prevalence of susceptibility and naturally acquired immunity, and to infer reporting rates. Our model accounted for transitions among susceptible *(S)*, exposed *(E)*, infectious (*I*), and recovered-immune *(R)* classes, *N = S+E+I+R*, assuming infectiousness begins, on average, σ^−1^ =17 days after exposure and lasts, on average, γ^−1^ =5 days (*54*). We partitioned the population across age classes (0-11m, 1-4y, 5-9y, 10-14y, 15-19y, 20-24y, 25-29y, 30-39y, 40-64y, ≥65y) corresponding to the ranges for reporting of aggregated case data (*55*), and defining the rate of aging from the *i* ^th^ class as *α*(*i*). For tractability we assumed a stable population, so that births into the youngest class were equal to deaths from the oldest class, i.e., μ(1) = α(10)*N*(10). The force of infection λ(*i*) is the rate of infection in the susceptible class. Taken together,

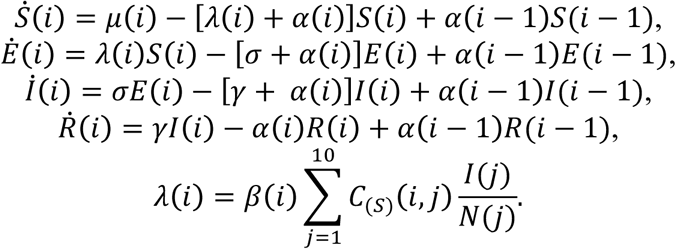

The force of infection *λ*(*i*) arises from age-structured social mixing in the context of differential age-specific prevalence of infection. We rescaled matrix elements *C*_(*S*)_(*i*, *j*) from diary data of daily, age-specific respiratory contacts from the UK (*56*) to accommodate the age distribution of the population, as modeled, with equal birth and death rates, and made elements symmetric

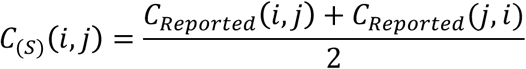

to correct for biased responses. We fitted *β*(*i*) for each age group to allow flexibility, for instance due to differences from contact patterns reported by survey participants, differences across ages in risk for infection given exposure (as may arise since young children are more likely to put their hands or objects in their mouths), and differential susceptibility across ages due to factors other than infection-derived or vaccine-derived immunity (for instance, maternal antibody-mediated protection in the first year of life).

Model parameters are listed with sources and definitions in Table S2. Replication code including parameter distributions and estimates of time-varying parameters is available at github.com/joelewnard/mumps.

**Table S2:**
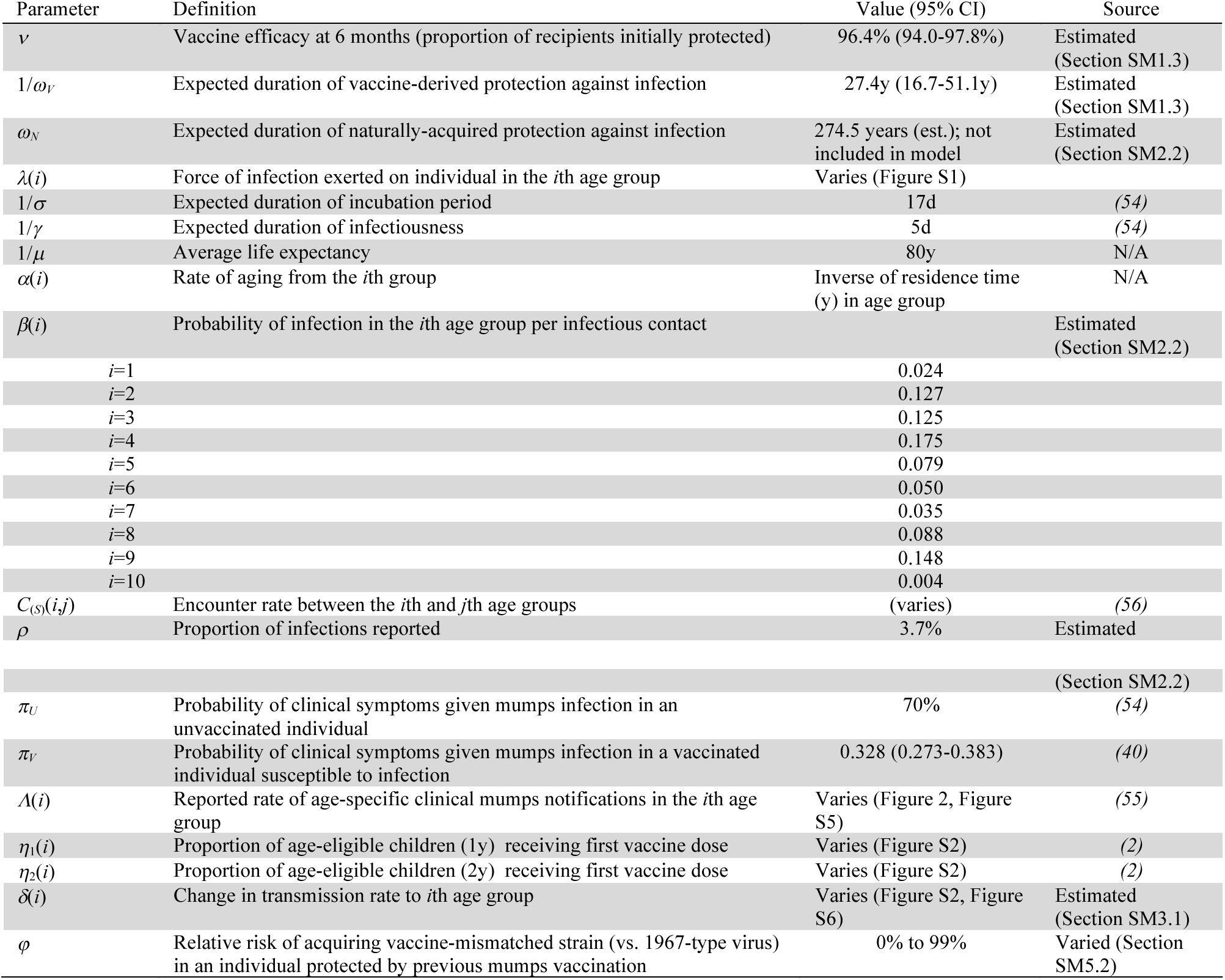
Model parameter definitions and values.

| Parameter | Definition | Value (95% CI) | Source |
| --- | --- | --- | --- |
| $\nu$ | Vaccine efficacy at 6 months (proportion of recipients initially protected) | 96.4% (94.0-97.8%) | Estimated (Section SM1.3) |
| $1/\omega_V$ | Expected duration of vaccine-derived protection against infection | 27.4y (16.7-51.1y) | Estimated (Section SM1.3) |
| $\omega_N$ | Expected duration of naturally-acquired protection against infection | 274.5 years (est.); not included in model | Estimated (Section SM2.2) |
| $\lambda(i)$ | Force of infection exerted on individual in the $i$th age group | Varies (Figure S1) | |
| $1/\sigma$ | Expected duration of incubation period | 17d | (54) |
| $1/\gamma$ | Expected duration of infectiousness | 5d | (54) |
| $1/\mu$ | Average life expectancy | 80y | N/A |
| $\alpha(i)$ | Rate of aging from the $i$th group | Inverse of residence time (y) in age group | N/A |
| $\beta(i)$ | Probability of infection in the $i$th age group per infectious contact | | Estimated (Section SM2.2) |
| $i=1$ | | 0.024 | |
| $i=2$ | | 0.127 | |
| $i=3$ | | 0.125 | |
| $i=4$ | | 0.175 | |
| $i=5$ | | 0.079 | |
| $i=6$ | | 0.050 | |
| $i=7$ | | 0.035 | |
| $i=8$ | | 0.088 | |
| $i=9$ | | 0.148 | |
| $i=10$ | | 0.004 | |
| $C_{(S)}(i, j)$ | Encounter rate between the $i$th and $j$th age groups | (varies) | (56) |
| $\rho$ | Proportion of infections reported | 3.7% | Estimated |
|  |  |  | (Section SM2.2) |
| $\pi_U$ | Probability of clinical symptoms given mumps infection in an unvaccinated individual | 70% | (54) |
| $\pi_V$ | Probability of clinical symptoms given mumps infection in a vaccinated individual susceptible to infection | 0.328 (0.273-0.383) | (40) |
| $\Lambda(i)$ | Reported rate of age-specific clinical mumps notifications in the $i$th age group | Varies (Figure 2, Figure S5) | (55) |
| $\eta_1(i)$ | Proportion of age-eligible children (1y) receiving first vaccine dose | Varies (Figure S2) | (2) |
| $\eta_2(i)$ | Proportion of age-eligible children (2y) receiving first vaccine dose | Varies (Figure S2) | (2) |
| $\delta(i)$ | Change in transmission rate to $i$th age group | Varies (Figure S2, Figure S6) | Estimated (Section SM3.1) |
| $\phi$ | Relative risk of acquiring vaccine-mismatched strain (vs. 1967-type virus) in an individual protected by previous mumps vaccination | 0% to 99% | Varied (Section SM5.2) |

#### SM2.2 Parameter estimation

We estimated parameters allowing the model to recapitulate reported age-specific incidence rates over the years 1960-1964 in the US (*57*). Simulating 150y of transmission to achieve steady-state conditions, and recording the force of infection and susceptible population at equilibrium, we used the Nelder Mead algorithm to perform weighted least-squares estimation

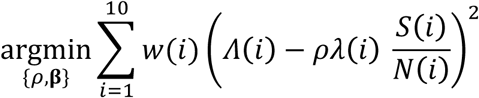

mapping reported age-specific incidence rates *Λ*(*i*) to model-predicted rates of infection, scaled by the proportion of infections reported (*ρ*). Consistent with an assumption that case frequencies are Poisson-distributed, we define weights *w*(*i,t*)=[1+*Λ*(*I,t*)]^−1^.

Our model-based estimate that *ρ*=3.7% closely corroborates the estimate of 4.0% (95%CI: 1.3– 8.0%) obtained from comparisons of surveillance data against community serosurveys (*58*). The basic reproductive number (*R*_0_=4.79) during the pre-vaccination era is the maximum eigenvalue of the matrix composed of elements

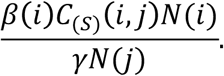

Providing a good fit to data, the model predicted 59% of individuals would acquire infection between the ages of 5 and 9, and that 89% would be infected by age 20 (**Figure S1**).

**Fig. S1:**
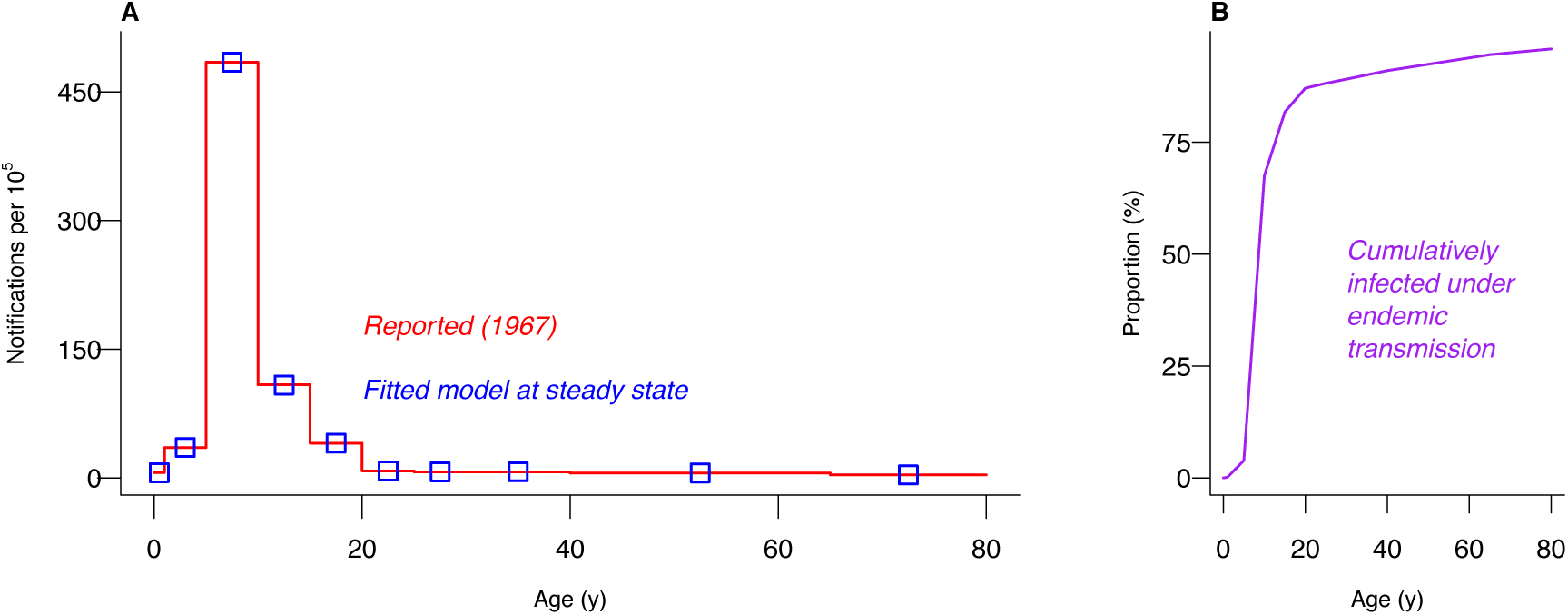
Fitted endemic transmission dynamics prior to vaccine rollout. We plot (**A**) reported and model-predicted case notification rates in the years 1960-1964 and (**B**) the corresponding proportion of previously infected and thus immune individuals under estimated transmission rates.

We also used the model to assess whether naturally-acquired immunity was likely to wane, assuming individuals exited the *R* class and re-entered the *S* class at a rate *ω*_N_. Our estimate that such waning would occur at an expected frequency of once per 274.5 years per person suggested it was unlikely, if extent, to be of epidemiologic importance. Moreover, we identified no improvement in model fit (defined as a lower value of the Akaike Information Criterion) when incorporating *ω*_N_ (AIC_0_ = −46.1, AIC_*ωN*_= −43.4).

### SM3. Modeling cohort-specific susceptibility to mumps under waning vaccine-derived protection

#### SM3.1 Relating case notification rates to transmission rates

We modified the transmission model described above (*section SM2.1*) to capture changes over time in the proportion of individuals susceptible to mumps (*S*), those with immunity acquired through natural infection (*I*), those with protection acquired from vaccination (*V*), and those who have experienced vaccine failure (*F*) due to an unsuccessful initial “take” (primary vaccine failure) or waning of initial protection (secondary vaccine failure). In the US, children have been recommended to receive vaccine doses at age 1 and between ages 4-6 before school entry. We model receipt of each dose in a proportion (**η**_1_ and **η**_2_) of individuals as they transition from the 0-11m to 1-4y age classes (first dose) and from the 1-4y to 5-9y age classes (second dose), assuming those who refuse the first dose do not seek out the second dose either. Defining *η*_1_(*i*)=0 for *i*>1 (corresponding to the 0-11m age group) and *η*_2_(*i*)=0 for *i*≠2 (corresponding to the 1-4y age group),

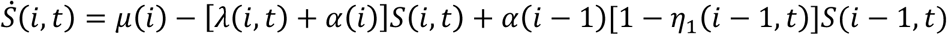

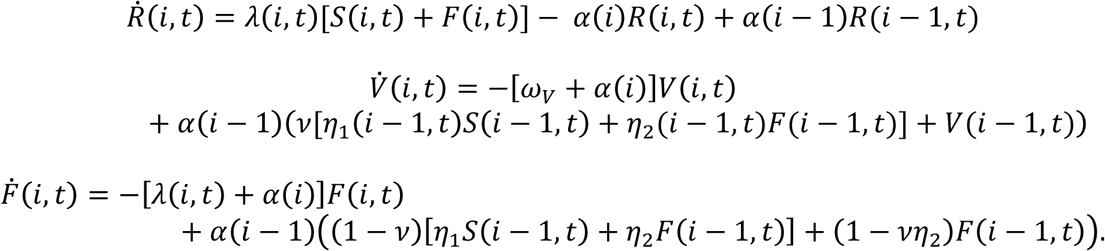

Numerous studies have shown that 70% (= *π*_*U*_) of mumps infections cause symptoms in previously susceptible, unvaccinated individuals (*54*, *59*, *60*), suggesting *ρ*/π_U_=5.3% of symptomatic infections are reported. Symptoms are estimated to occur in 27.0% to 38.6% of vaccinated individuals, suggesting 44.5% to 61.4% partial protection against symptoms persists even when vaccination fails to prevent infection due to primary or secondary failure (*40*). As this range was calculated assuming differing serological cutoffs for infection in the absence of a gold standard, we took *π*_*v*_ ∼ Unif(0.270,0.386). Such estimates of vaccine effectiveness against symptoms given infection are consistent with evidence of moderate protection against complications (including endpoints such as orchitis, meningitis, viruria, pancreatitis, and hospitalization) in studies of mumps infections among vaccinated and unvaccinated persons (*61*–*63*).

We back-calculated the cumulative force of infection to which individuals in each age group were exposed each year by updating the *S, R, V*, and *F* classes, thereby relating transmission to case notification data according to the expected proportion of infections causing symptoms and being reported:

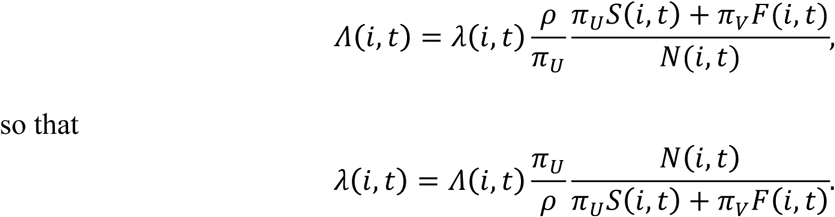

We obtained time series of the mean annual force of infection for each of 1,000 draws from the independent distributions of {*v*, ***Ŷ***^*^} and *π*_v_. As a sensitivity analysis, we repeated the procedure allowing the proportion of infections reported to vary over time (*section SM3.2*).

In addition to providing estimates of the population susceptible to mumps each year since vaccine rollout, this approach allowed us to assess changes over time in transmission rates that may result from changes in demography and contact patterns, behavior, health status, and response efforts aiming to limit transmission. We quantify changes in the rate of mumps acquisition per infectious exposure over time as

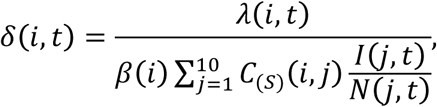

obtaining estimates of *R*_0_(*t*) as the maximum eigenvalues of the matrix composed of elements

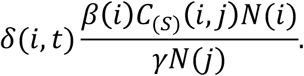

Declining values of *R*_0_(*t*) over time from 4.79 in the pre-vaccination era to 2.41 (1.85-3.38) as of 2016 indicate that reductions in mumps incidence after vaccine licensure have been attributable to changes in transmission dynamics beyond individual and herd protection conferred by the vaccine (**Figure S2**). This finding persisted in models assuming annual reductions in reporting (*section SM3.2*).

**Fig. S2:**
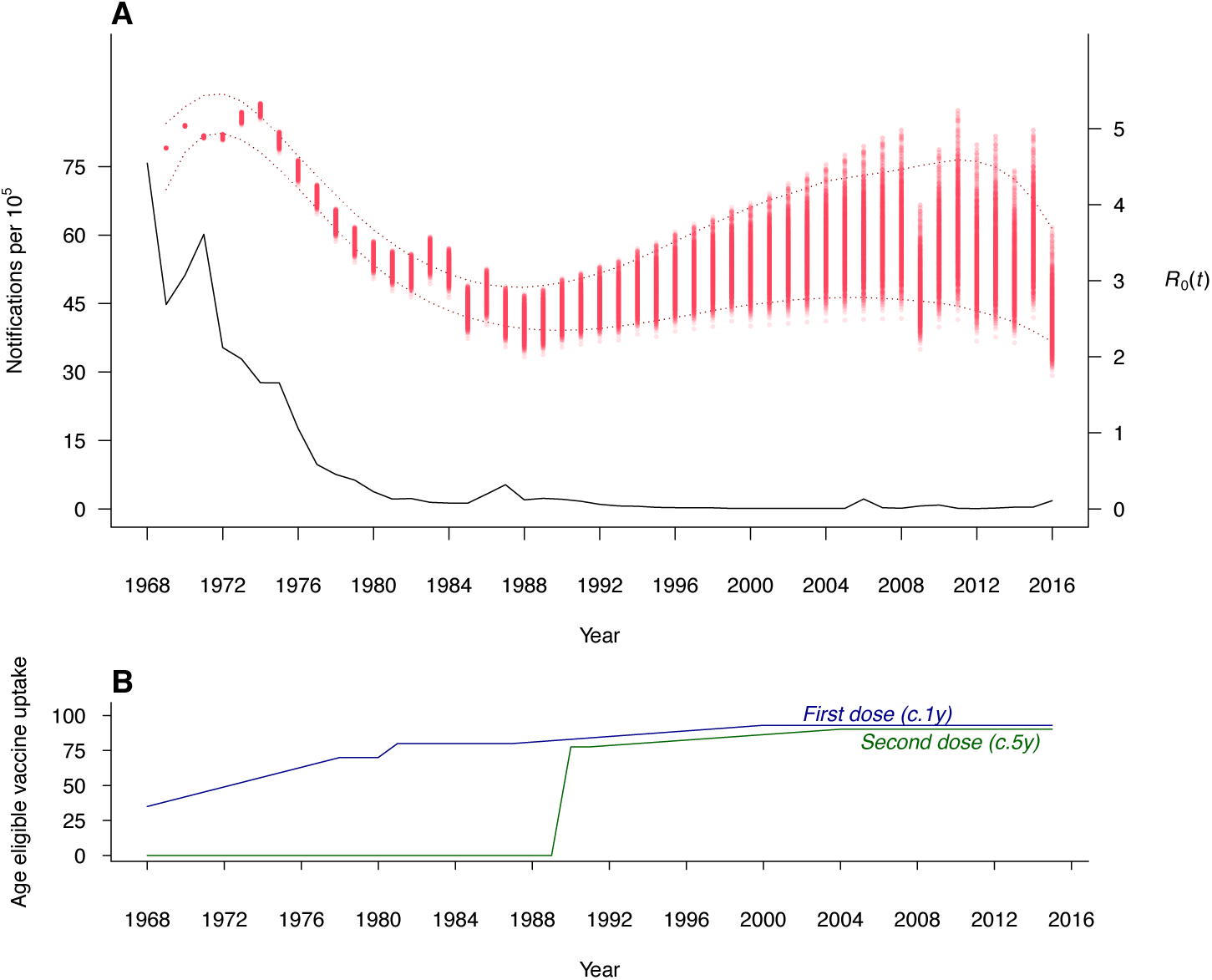
Reductions in mumps notifications correspond with increases in vaccine coverage and a declining basic reproductive number. We plot (**A**) reported annual incidence rates in the US against our estimates of *R*_0_(*t*) and (**B**) age-eligible annual vaccine uptake for the first dose (proportion of children receiving at age 1y) and second dose (proportion of children receiving at age 5y), using data from (*2*). Dotted lines delineate 95% confidence intervals around estimates of *R*_0_(*t*) as a continuous function of time, fitted individually to evaluations of the conditional distribution of *δ*(*i,t*) |{*v, ω* _*V*_, *π*_*V*_}. For each replicate, polynomial terms were added until no improvement was evident in values of the Bayesian Information Criterion.

Our inference of reduced transmission potential during the second half of the twentieth century is consistent with declines in the burden of other pediatric infectious diseases including pneumonia (*64*), rotavirus diarrhea (*65*), and central nervous system infections (*66*), which preceded licensure of vaccines against the causative agents *(Streptococcus pneumoniae*, rotavirus, and *Haemophilus influenzae* B and varicella, respectively). Many factors might contribute to these overall reductions in pediatric infectious disease burden. These changes correlate with the prevention of measles-induced immune suppression through widespread measles vaccination (*67*); with improvements in nutrition and health status suggested by declining prevalence of underweight and anemia in childhood (*68*, *69*); and with declining per-capita birth rates that have contributed to changes in the dynamics of measles, pertussis, and other childhood diseases in tandem with vaccination over the same period (*70*–*72*. In addition, implementing enhanced interventions in response to mumps cases amid vaccine rollout could be a factor in reduced transmission potential. Whereas mumps infections were once routine for children, outbreaks in the decades following vaccine licensure have prompted epidemiologic investigations and containment measures including active surveillance and isolation of cases (*30*, *31*, *73*, *74*). Evidence that vaccinated and unvaccinated persons who acquire infection have equal viral loads in saliva and urine suggests that diminished capacity of vaccinated, infected persons to transmit is not a factor in the observed decline in transmission per infected person (*62*).

To accompany our analysis of changes in age-specific incidence and susceptibility, we also plotted changes in cohort-specific incidence over time (**Figure S3**). We determined the distribution of birth years among annual reported cases by defining annual incidence as binomially distributed within age groups, with *n* equal to the population of each age and *p* the cases per capita per year; we pooled these distributions to calculate the median and interquartile range of birth years among reported cases. Our outcomes illustrate that cases reported during the outbreaks occurring in the late 1980s and early 1990s arose mostly among individuals recommended to receive one vaccine dose, who were exposed to reduced rates of mumps transmission in childhood compared to earlier cohorts. Rather than reflecting a continuation of cases within this under-immunized cohort, the resurgence of mumps from 2006 to the present has predominantly affected individuals recommended to receive two vaccine doses, consistent with reports from outbreak investigations (*30, 46, 47, 53*).

**Fig. S3:**
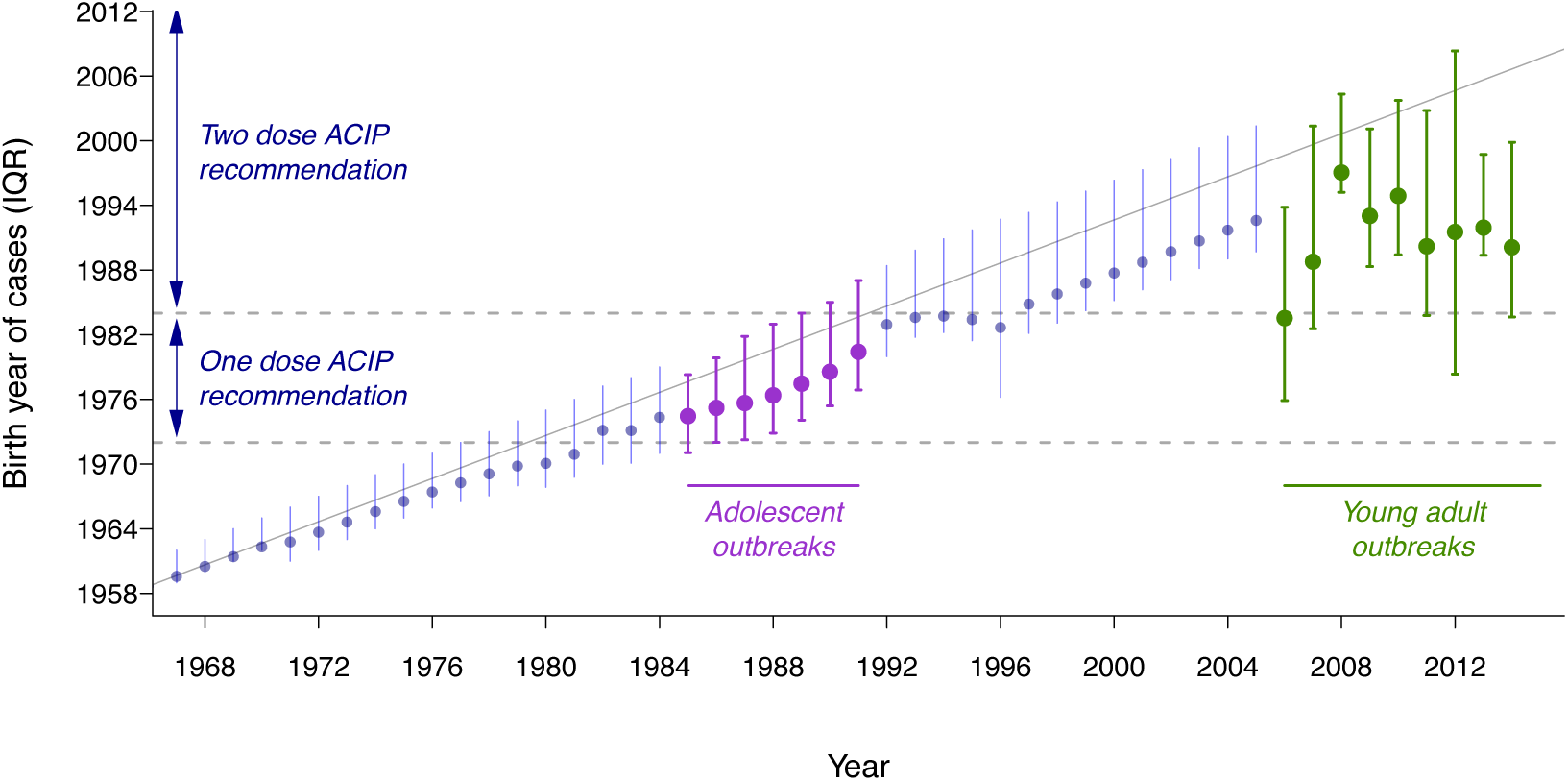
Birth cohorts accounting for reported cases over time. Illustrating the birth years of reported cases demonstrates that outbreaks following vaccine introduction have affected distinct cohorts. Points and vertical lines indicate medians and interquartile ranges (IQRs) of the birth year of reported cases; the diagonal grey line indicates the expected median age of cases under a continuation of endemic transmission from 1967 forward. Horizontal, dashed lines indicate the birth years associated with one-dose and two-dose vaccination ACIP recommendations, not accounting for state differences in catch-up effort.

#### SM3.2 Sensitivity analysis under time-varying reporting

We assessed whether inferred levels of protection and changes in transmission dynamics could be subject to bias resulting from lower reporting of mumps infections after vaccine rollout. Suggested by our finding of lower *R*_0_ values over time, such declines may have arisen in the context of lower public and clinical familiarity with mumps due to reductions in incidence, or if the occurrence of cases outside their historical range of 5-9y impacted detection.

We identify no appreciable changes (>15% departure from original estimate) in age-specific susceptibility when considering 1% and 2% annual reductions in the probability (*ρ*/*π*_*U*_) for symptomatic cases to be reported—corresponding to 39% and 63% overall reductions in reporting of symptomatic cases, respectively, between 1967 and 2016 (**Figure S4**). As an increasing proportion of population immunity has been attributable to mumps vaccination rather than natural mumps infection, the impact of these changes on absolute estimates of the number of people immune to mumps infection declines toward 2016.

**Fig. S4:**
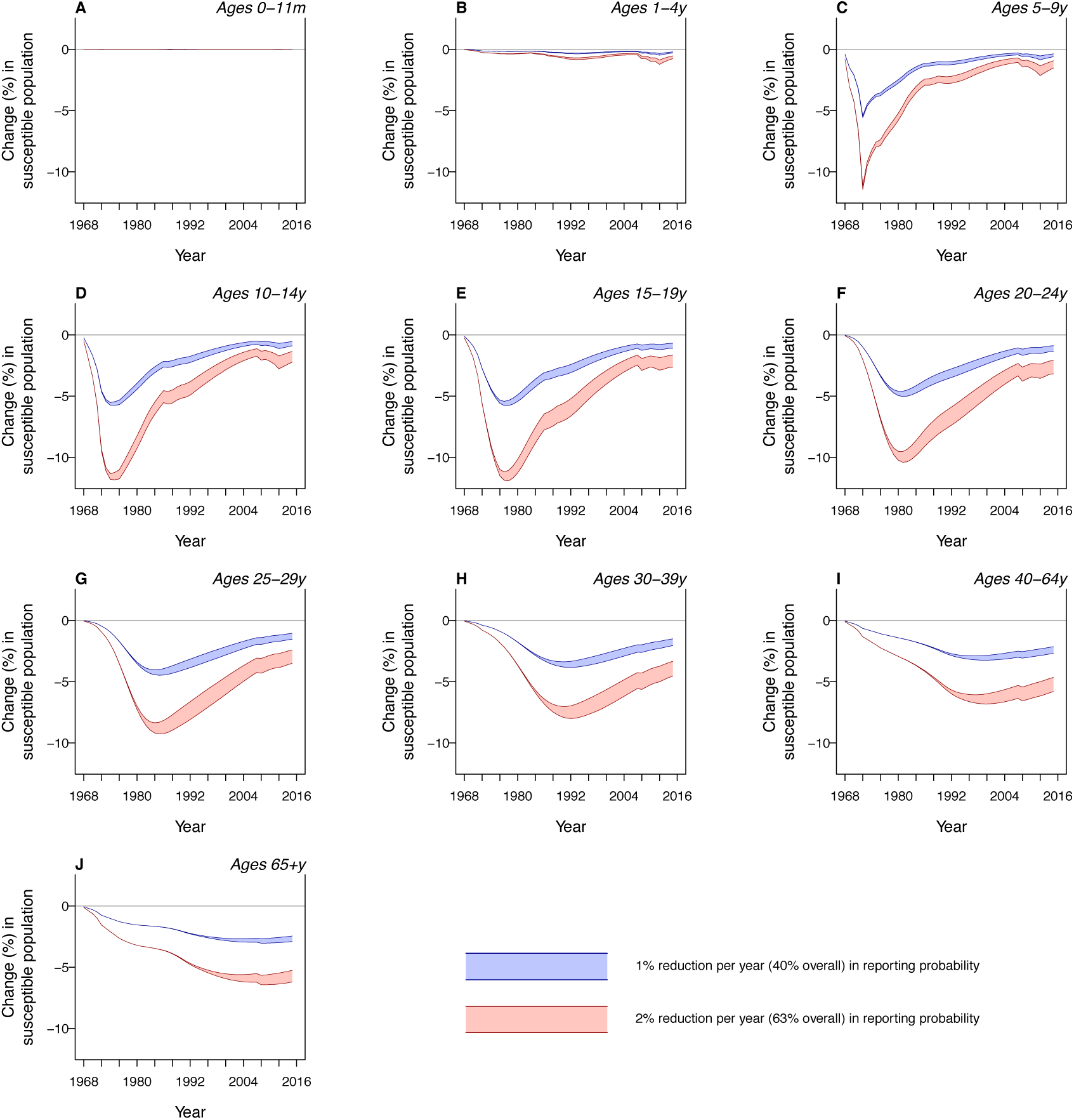
Changes in estimates of the susceptible population under scenarios of declining reporting. An assumption of declining reporting rates results in <15% reductions in estimates of age-specific prevalence of susceptibility. The impact is lower at later time points (5% or lower across ages <65y as of 2016), reflecting the low degree of transmission in recent decades and greater impact of vaccination on population susceptibility. Shaded areas delineate 95% confidence intervals.

Moreover, estimates of *R*_0_ suggest our inferences of declining transmission potential are robust to potential declines in reporting (**Figure S5**). Compared to an original estimate of 2.41 (1.85-3.38) as of 2016 assuming constant reporting, our estimates of 2.43 (1.73-3.28) under 1% annual decreases in reporting and 2.43 (1.74-3.28) under 2% annual decreases demonstrate that this inference is not sensitive to changes in reporting.

**Fig. S5:**
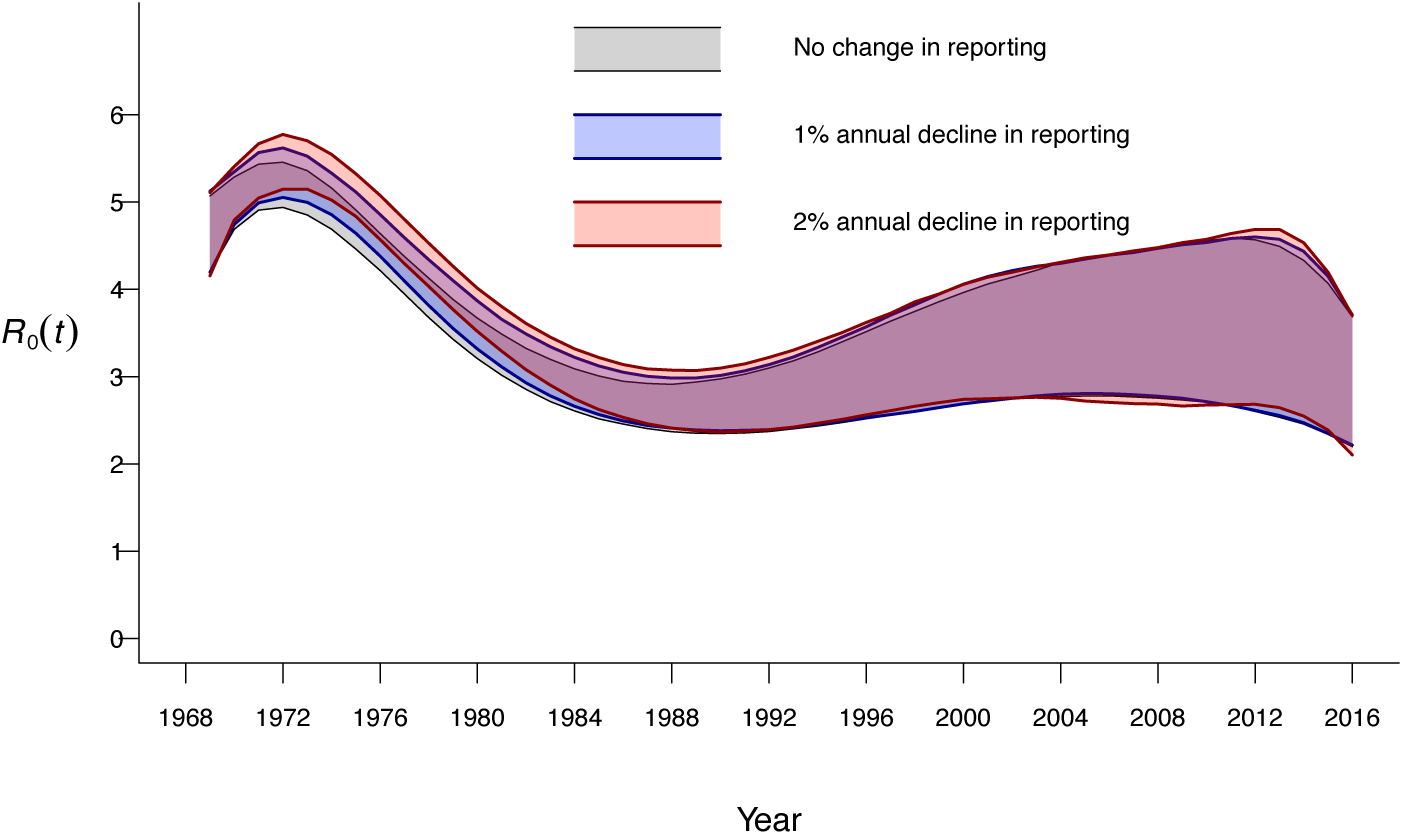
Changes in estimates of *R*_0_ over time under scenarios of declining reporting. An assumption of declining reporting rates has minimal bearing on estimates of the basic reproductive number. Shaded areas delineate 95% confidence intervals. Dotted lines delineate 95% confidence intervals around estimates of *R*_0_(*t*) as a continuous function of time, fitted individually to evaluations of the conditional distribution of *δ*(*i,t*) |{*v, ω* _V_, *π*_*V*_}. For each replicate, polynomial terms were added until no improvement was evident in values of the Bayesian Information Criterion.

#### SM3.4 Interpolating age-specific annual incidence from case notification data

Our analysis uses annual, age-specific incidence rates organized into strata of ages 0-11m, 1-4y, 5-9y, 10-14y, 15-19y, 20-24y, 25-29y, 30-39y, 40-64y, and ≥65y. While these strata were defined based on the usual aggregation of case notifications by the Centers for Disease Control and Prevention (*55*), aggregations differed by year. When aggregations occurred at smaller (e.g., one year of age) intervals, we calculated the stratum-wide incidence rate as a population-weighted mean across age-specific incidence rates within the stratum. When aggregations crossed age strata (e.g., incidence reported for ages 20-29y, rather than 20-24y and 25-29y), we used linear interpolation to estimate incidence in the stratum of interest based on the ratio of incidence in the stratum of interest to incidence across the aggregated strata in the years immediately preceding and following. For instance, if rates were reported for ages 20-29 in year *t*, we computed incidence at ages 20-24y assuming

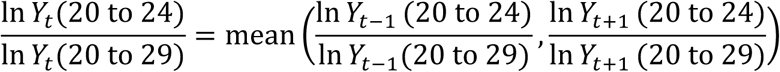

Overall annual incidence was available from each year, enabling us to use the same linear interpolation approach to estimate age-specific rates when age-specific notifications were aggregated over multi-annual periods (up to 5y long in the data). We illustrate the aggregation of reported data by year and by age group (**Figure S6**) together with estimated age-specific rates. Replication files are available in the github repository referenced in the main text.

**Fig. S6:**
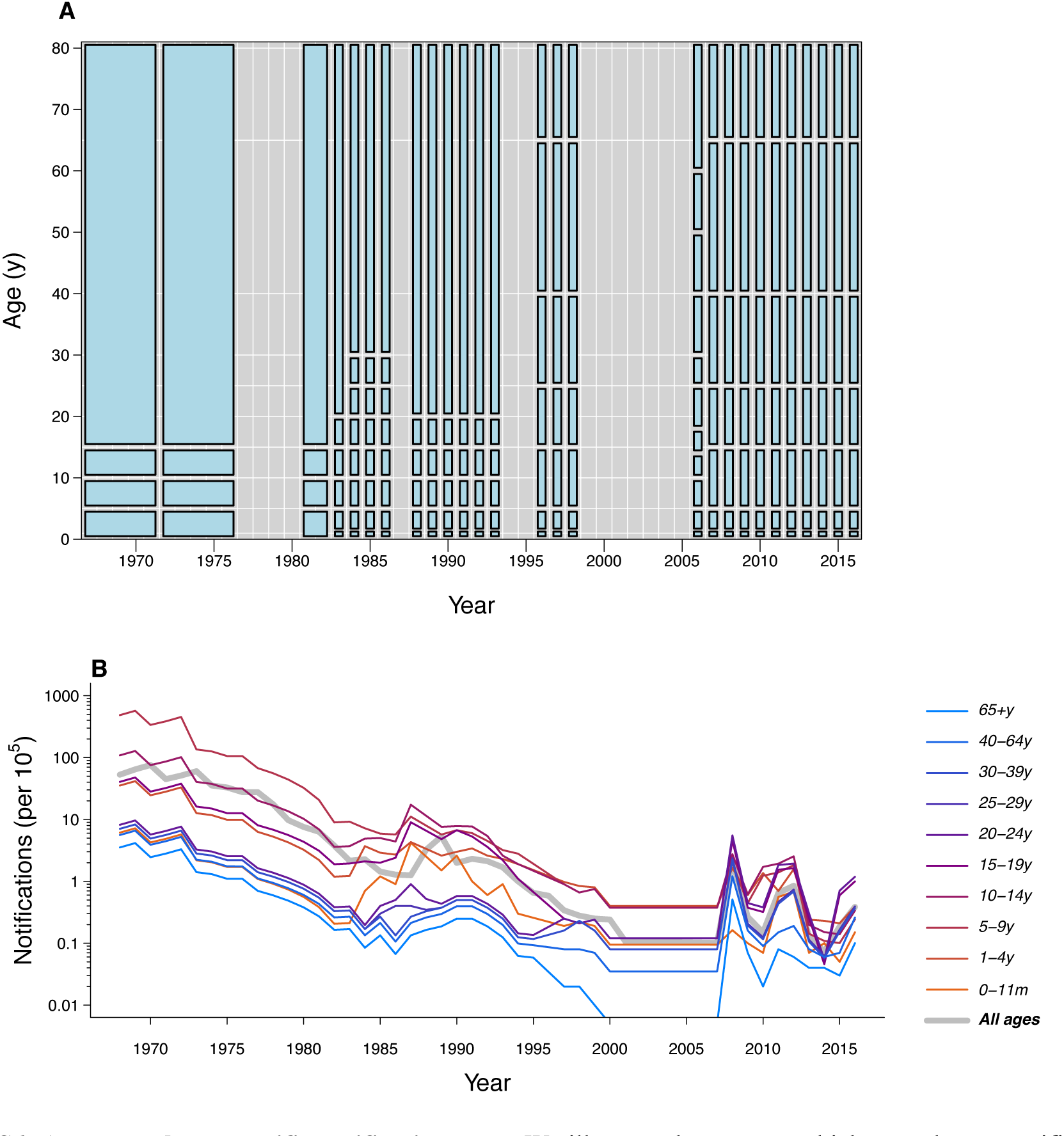
Aggregated age-specific notification rates. We illustrate the strata at which annual age-specific notification rates were reported as bins (**A**) together with estimated incidence rates by age group (**B**). Reporting of mumps cases was discontinued from 1999 to 2005 (*55*). We digitized aggregated incidence over this period from Figure 1 in (*2*) assuming constant proportions of cases by age; this assumption had little impact since overall notification rates were low (<1 per 100,000) over this period.

### SM4. Modeling cohort-specific susceptibility to mumps assuming no waning of vaccine-derived protection

As an alternative to the hypothesized role of waning protection, the emergence of mumps virus strains escaping vaccine-derived protection has received attention as a potential explanation for the resurgence of mumps since 2006 (*9*, *12*) and reduced vaccine effectiveness in recent outbreaks (*14, 48, 50, 53*). To provide a basis for simulating transmission dynamics under this scenario (*section SM5.2*), we solved for the force of infection *λ*(*i,t*), updated the populations of the *S*, *R*, *V*, and *F* states, and estimated changes in transmission *δ*(*i,t*) using the approach described in previously (*section SM3.1*), this time assuming waning of vaccine-derived immunity
was not the explanation for continued transmission (*ω*_*V*_=0); naturally, longer-lasting vaccine protection leads to higher estimated proportions of individuals protected against vaccine-type (pre-2006) lineages (**Figure S7**). In contrast to our estimates of declining *R*_0_ values over time in the model with waning protection (*SM3.1*), we infer that the basic reproductive number would have increased to 8.20 (7.46, 9.09) in order to sustain observed transmission at levels of immunity expected in the population expected without waning of vaccine derived protection.

**Fig. S7:**
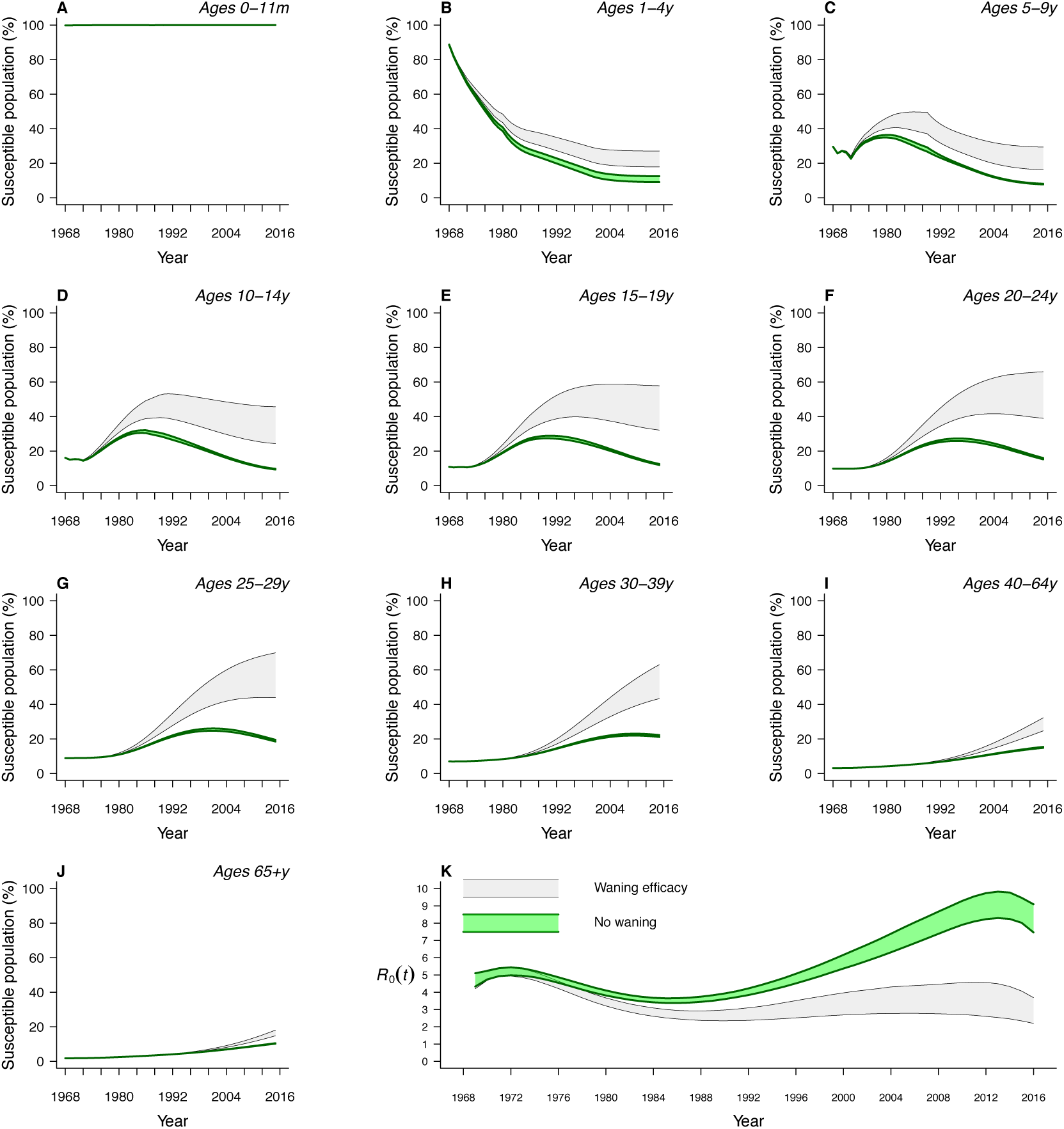
Estimates of population susceptibility and *R*_0_(*t*) under an assumption of time-invariant protection. (**A**–**J**) Under the assumption that protection against mumps does not wane with time since vaccination, the susceptible population is estimated to have decreased at ages <15y and to have increased to a lesser extent at older ages. Transient increases in susceptibility at ages 5-9y, 10-14y, 15-19y, and 20-24y reflect reduced incidence and incomplete vaccine uptake during the decades immediately following vaccine rollout. (**K**) We infer that, for transmission to be sustained at reported levels, *R*_0_ would have increased nearly two-fold to 8.20 (7.46, 9.09) as of 2016. Shaded areas delineate 95% confidence intervals.

### SM5. Simulating mumps re-emergence under situations of waning vaccine-derived protection and vaccine escape

#### SM5.1 Simulating mumps transmission dynamics under waning vaccine-derived protection

We next sought to understand whether the abrupt resurgence in mumps notifications in the US in 2006 was more consistent with dynamics predicted under a situation of vaccine waning or of vaccine escape. Under the scenario of waning protection, we extend the model presented in SM2.1 to include vaccination, as described in SM3.1, obtaining the system of equations

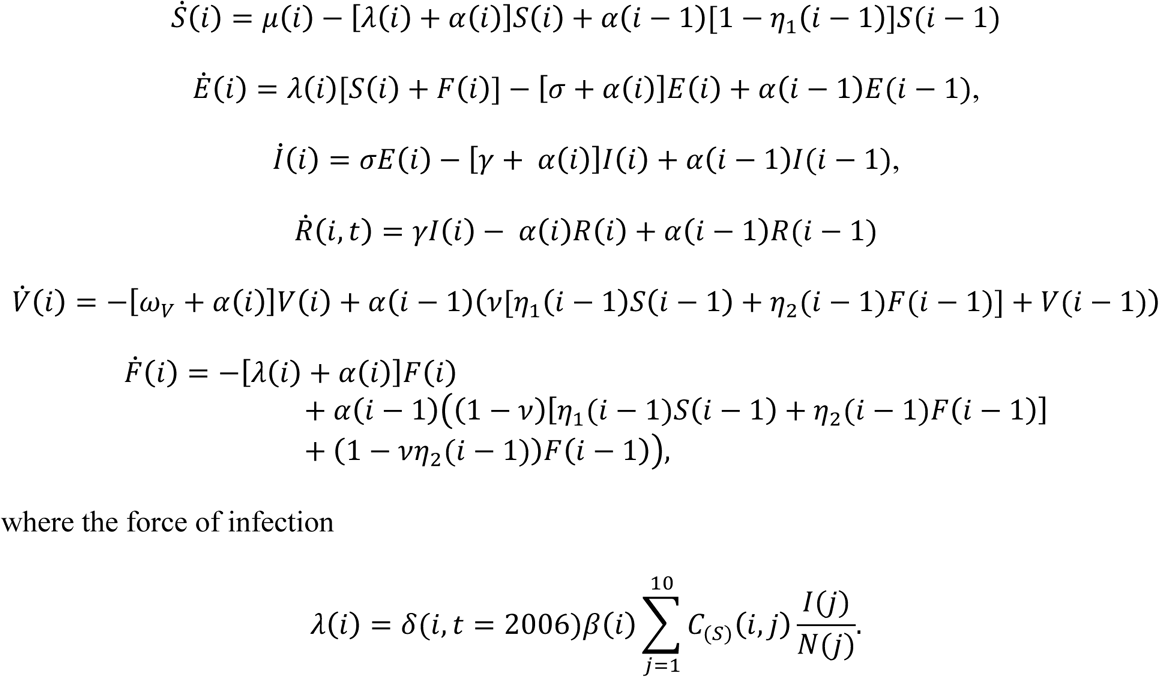

We initialize the model with estimates of the population across *S*, *R*, *V*, and *F* states as of 2006, assuming introductions at randomly-drawn frequencies below expected endemic prevalence. We simulate dynamics in a hypothetical population of 1 million for one year using the Gillespie (stochastic simulation) algorithm, assuming exponentially-distributed event times (*75*). We assume infections among susceptible persons are detected with probability *ρ*, while infections among vaccinated persons are detected with probability *ρπ*_*V*_/π_*U*_ due to partial protection against symptoms.

#### SM5.2 Simulating mumps transmission dynamics under vaccine escape

We modified the model to assess the potential transmission dynamics of a mumps virus strain with a partial ability to escape vaccine-derived immune protection. Defining *φ* (0<*φ*<1) as the probability for the introduced strain to infect an exposed individual protected by vaccine-derived immunity, we modify the differential equations above (section *SM5.1*) as follows:

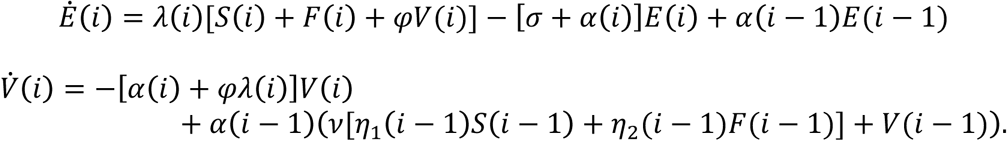

Using transmission parameters *δ*_*ωv*_=0{*i, t*) inferred under the assumption that vaccine-derived protection does not wane, we update the force of infection as

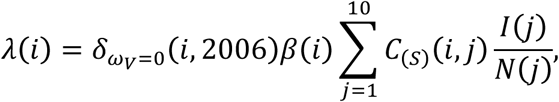

simulating with *ω*_*V*_=0. We further assume that although vaccinated individuals may be at risk for acquiring the escape strain, naturally immune individuals are protected. We arrive at this consideration based on evidence of stronger immune responses following natural infection in comparison to vaccination: average anti-mumps neutralizing antibody titers are approximately 16-fold higher in naturally-infected children compared mumps-vaccinated children (*18*), consistent with observations in measles (*76*, *77*) and rubella (*78*), among other viral infections of childhood (*36*, *79*). At adequate concentrations, such antibodies are expected to neutralize heterologous strains effectively (*25*, *26*).

#### SM5.3 Comparing predicted dynamics from stochastic simulations

To determine which scenario was more concordant with observed cases, we compared model-predicted age-specific and overall incidence rates against reported rates, as well as the predicted and reported median age of infection. To obtain the predicted median age of infection, we constructed an *in silico* pseudo-population corresponding to the US age distribution as of 2006 (ages 0, 1, 2, …, 80y) and drew multinomial samples of infection ages according to model-predicted total age-specific incidence over one year, calculating the median at each iteration.

#### SM5.4 Age of infection and vaccine escape

Our analysis suggested that endemic transmission of virus strains escaping vaccine-induced immunity may be expected to result in cases following a younger (e.g., 5-18y) age distribution. Most recent outbreaks in the US have centered among young adults. However, a large outbreak in Arkansas in 2016-17 took place primarily in children ages 5-17y (*80*). While this outbreak was connected to a larger epidemic involving adults in the Marshall Islands, it is unclear whether vaccine escape played an additional role contributing to the occurrence of infections at young ages in Arkansas. Such circumstances motivate genomic epidemiological studies which can assess whether outbreaks affecting children involve transmission of distinct mumps virus strains or are connected to transmission among susceptible young adults whose immunity has waned, to low rates of vaccination in the population, or to other factors.

### SM6. Anticipating third-dose impact

#### SM6.1 Expected protection across ages

The continued aging of individuals who were exposed to high rates of mumps transmission in childhood prior to the 1980s suggests the US population will, in the future, depend largely on vaccine-derived direct and indirect protection against mumps. We estimated levels of age-specific protection attainable under the current vaccine dosing schedule, as well as an extended three-dose schedule, to assess the potential for eliminating transmission through vaccination. Simplifying the model from section SM3.1 to account for protection resulting only from immunization,

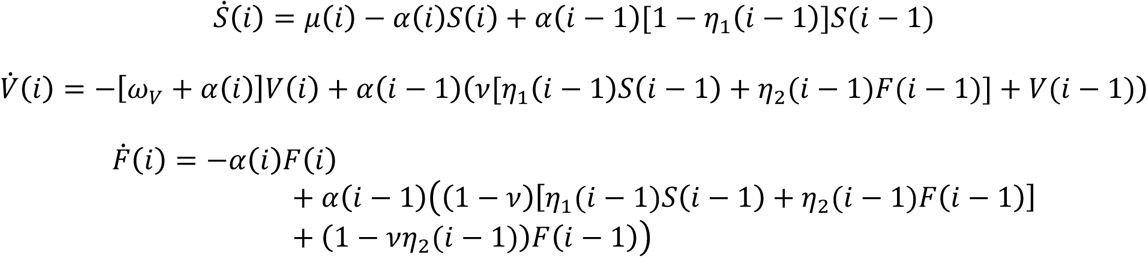

under the current approach with doses administered at ages 1y and 5y, approximately. With an added third dose administered between ages 15-19y at a rate *r*_3_(*i*),

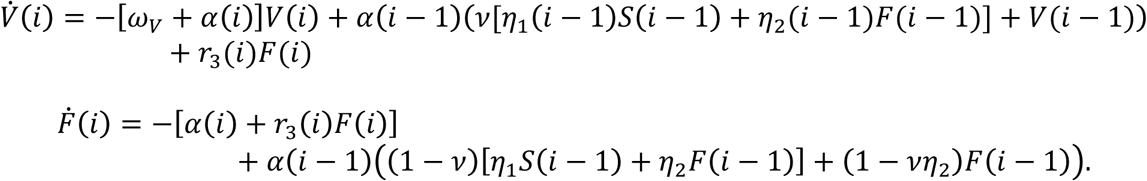

We also considered scenarios including boosters at 10y and 20y frequencies for individuals ages 20y and older, included in the above equations via the rate *r*_3_(*i*). We assumed two scenarios for uptake of these added doses. As a pessimistic case, we assumed that third-dose and booster coverage would resemble the 56% coverage of tetanus-diptheria (Td) booster doses in the US adult population (*81*). To obtain this coverage level in the population level as a whole, we modeled uptake among 62% of individuals who completed the recommended primary series. As an optimistic case, we assumed the probability for second-dose recipients to obtain a third dose (and for individuals to continue receiving subsequent booster doses) would equate the probability for first-dose recipients to obtain a second dose (97%), resulting in 88% overall third-dose coverage.

#### SM6.2 Evaluating effective reproductive numbers under extended vaccine dosing schedules

To understand the impact of the vaccine schedules under consideration on transmission dynamics, we calculated the effective reproductive number *R*_*E*_(*t*) under steady-state distributions of age-specific susceptibility as the maximum eigenvalue of the matrix with entries

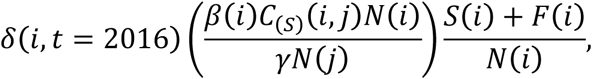

using estimates of *δ*(*i,t*) as of 2016 to account for changes in transmission rates.

To determine the threshold level of vaccine protection against emerging strains allowing *R*_*E*_=1, we defined *R*_*E*_ of an emerging strain as the maximum eigenvalue of a matrix with entries
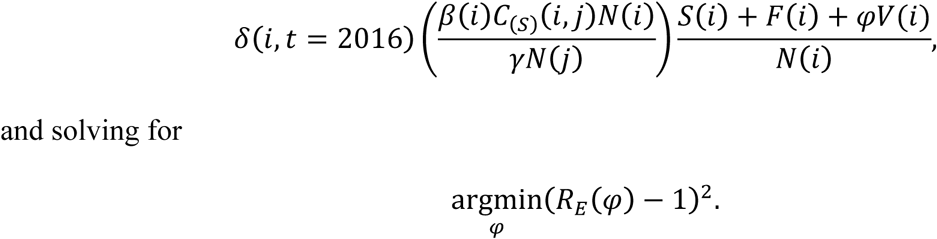

